# Dendritic calcium signals in rhesus macaque motor cortex drive an optical brain-computer interface

**DOI:** 10.1101/780486

**Authors:** Eric M. Trautmann, Daniel J. O’Shea, Xulu Sun, James H. Marshel, Ailey Crow, Brian Hsueh, Sam Vesuna, Lucas Cofer, Gergő Bohner, Will Allen, Isaac Kauvar, Sean Quirin, Matthew MacDougall, Yuzhi Chen, Matthew P. Whitmire, Charu Ramakrishnan, Maneesh Sahani, Eyal Seidemann, Stephen I. Ryu, Karl Deisseroth, Krishna V. Shenoy

**Affiliations:** Neurosciences Graduate Program, Stanford University, Stanford, CA 94305, USA; Department of Electrical Engineering, Stanford University, Stanford, CA 94305, USA; Department of Biology, Stanford University, Stanford, CA 94305, USA; Department of Bioengineering, Stanford University, Stanford, CA 94305, USA; Gatsby Computational Neuroscience Unit, University College London, London W1T 4JG, UK; Department of Neurosurgery, Stanford University, Stanford, CA 94305, USA; Center for Perceptual Systems, University of Texas, Austin, TX 78705, USA; Department of Psychology, University of Texas, Austin, TX 78705, USA; Department of Neuroscience, University of Texas, Austin, TX 78705, USA; Department of Neurosurgery, Palo Alto Medical Foundation, Palo Alto, CA 94301, USA; Department of Psychiatry and Behavioral Science, Stanford University, Stanford, CA 94305, USA; Howard Hughes Medical Institute, Stanford University, Stanford, CA 94305, USA; Wu Tsai Neuroscience Institute, Stanford University, Stanford, CA 94305, USA; Department of Neurobiology, Stanford University, Stanford, CA 94305, USA

## Abstract

Calcium imaging has rapidly developed into a powerful tool for recording from large populations of neurons *in vivo*. Imaging in rhesus macaque motor cortex can enable the discovery of new principles of motor cortical function and can inform the design of next generation brain-computer interfaces (BCIs). Surface two-photon (2P) imaging, however, cannot presently access somatic calcium signals of neurons from all layers of macaque motor cortex due to photon scattering. Here, we demonstrate an implant and imaging system capable of chronic, motion-stabilized two-photon (2P) imaging of calcium signals from in macaques engaged in a motor task. By imaging apical dendrites, some of which originated from deep layer 5 neurons, as as well as superficial cell bodies, we achieved optical access to large populations of deep and superficial cortical neurons across dorsal premotor (PMd) and gyral primary motor (M1) cortices. Dendritic signals from individual neurons displayed tuning for different directions of arm movement, which was stable across many weeks. Combining several technical advances, we developed an optical BCI (oBCI) driven by these dendritic signals and successfully decoded movement direction online. By fusing 2P functional imaging with CLARITY volumetric imaging, we verify that an imaged dendrite, which contributed to oBCI decoding, originated from a putative Betz cell in motor cortical layer 5. This approach establishes new opportunities for studying motor control and designing BCIs.

## Introduction

Understanding the mechanisms by which populations of neurons give rise to behavior is a primary goal of systems neuroscience, and a foundational step for developing brain-computer interfaces (BCIs) for treating people with neurological injury and disease. BCIs seek to restore lost function by decoding neural activity from the brain in real time to control a medical device such as a prosthetic arm or computer interface^1, 2^. Investigations using rhesus macaques (*Macaca mulatta*) have contributed substantial progress towards developing clinically-viable BCIs by exploring decoding algorithms and system designs and by advancing our basic scientific understanding of the motor system^3–13^.

Typically, such experiments in monkeys use intracortical multi-electrode arrays, which are nearly identical to those approved for use in humans (e.g. so-called “Utah arrays”), to readily translate scientific and technical achievements from pre-clinical research in monkeys to improvements in clinical BCIs. For some aspects of BCI implementation and translation, such as studying biocompatibility, stability and longevity, using the same implantable sensor is of central importance. But for other aspects of BCI experimentation, the central goal is the scientific study of neural population dynamics underlying the control of arm movements in order to obtain fundamental understanding that can inform the design of future high-performance BCI systems^14^. For this goal, the tremendous recent expansion of tools for measuring activity from large populations of neurons in non-human model organisms could provide a fertile experimental landscape for exploring the design of next-generation BCIs. Here, we describe such an approach by using two photon (2P) imaging in macaque motor cortex to implement an optical BCI (oBCI).

In recent years, techniques for imaging the activity of large populations of neurons have improved rapidly. Imaging calcium dynamics using GCaMP is a popular and widely used modality and has proved successful for measuring primary visual cortex in macaques^15–19^ and in marmosets^20, 21^, as well as primary motor cortex in marmosets^22^. While calcium imaging methods do not yet recover the finely time-resolved spiking activity that electrophysiology does, they capture a complementary view of neural population activity by contextualizing activity within a dense, spatially localized, and genetically annotated map of the neural tissue. Optical methods can also readily access large neural populations^23, 24^, and can access additional neurons or brain areas simply by translating the objective lens or adjusting the scan pattern. An oBCI would thereby be particularly well suited for enabling researchers to explore the design space of how best to measure from neural populations (which area(s) to record from, how many neurons are needed, electrode density and distribution, etc). The knowledge gained can help set the design specifications for future electrode-array based BCIs. Using optical techniques, it is possible to dissociate the limitations of present-day electrode arrays, which are surgically implanted in a fixed location, from the fundamental study of neural population activity critical to the design of next-generation BCIs.

In this study, we developed an oBCI that operated by decoding neural population signals in real time at single-cell resolution in macaque motor cortex, with the capability of accessing signals from neurons in both superficial and deep layers. Neurons throughout the cortical lamina display tuning for different movements, making it desirable to access signals from both superficial and deep layers^25^. However, due to the light-scattering properties of brain tissue, it is not currently possible to image somatic calcium signals of deep layer 5 neurons that serve as the primary motor cortical output of movement signals to subcortical motor circuits and the spinal cord. Reliably imaging neuronal somas at this depth would require fundamental advances in physics and imaging technology or surgical implantation of a large (>1 mm diameter) penetrating lens^26, 27^. Implanting such a lens, however, has drawbacks: it requires lesioning and occasionally removing a large volume of tissue immediately adjacent to the imaged tissue region, potentially severing input and output connections to the imaged tissue. It is unclear the extent to which this tissue disruption impacts the neural population dynamics of the tissue under study. In addition, to image a new population of neurons in a different field of view with a penetrating lens requires a new implantation and additional tissue disruption. This is of particular concern if penetrations are performed in multiple locations to sample different neural populations, which realistically constrains this approach to one or a small handful of penetrations per subject. In addition, a penetrating lens, if fixed to the skull, is likely to further damage surrounding tissue if the brain pulsates due to changes in intracranial pressure, making it unclear how long such an implant would last in practice.

Here we developed a fundamentally different approach to record signals from neurons in deep layers, in addition to accessing neurons in superficial layers. This is of high value because neurons across all layers potentially generate signals relevant to understanding motor control and for driving BCI decoders^25, 28^. Cortical neurons, particularly layer 5 output neurons, extend apical dendrites to the most superficial layer of cortex. These apical dendrites have been previously shown to represent behaviorally relevant information using calcium imaging in mice^29–32^, presenting an opportunity to record signals originating from deep neurons by imaging their superficial compartments, shown schematically in Fig. 1. This possibility is particularly intriguing in light of recent work demonstrating that dendritic calcium transients are highly correlated with somatic activity in layer 5 neurons^33^. Nevertheless, to our knowledge, these superficial signals have not been specifically recorded using electrical or optical methods for either basic science or BCI applications in macaques or other nonhuman primates (NHPs); thus, it is unknown whether these superficial non-somatic signals could drive a BCI and provide insight into neural population dynamics.

**Figure 1.**
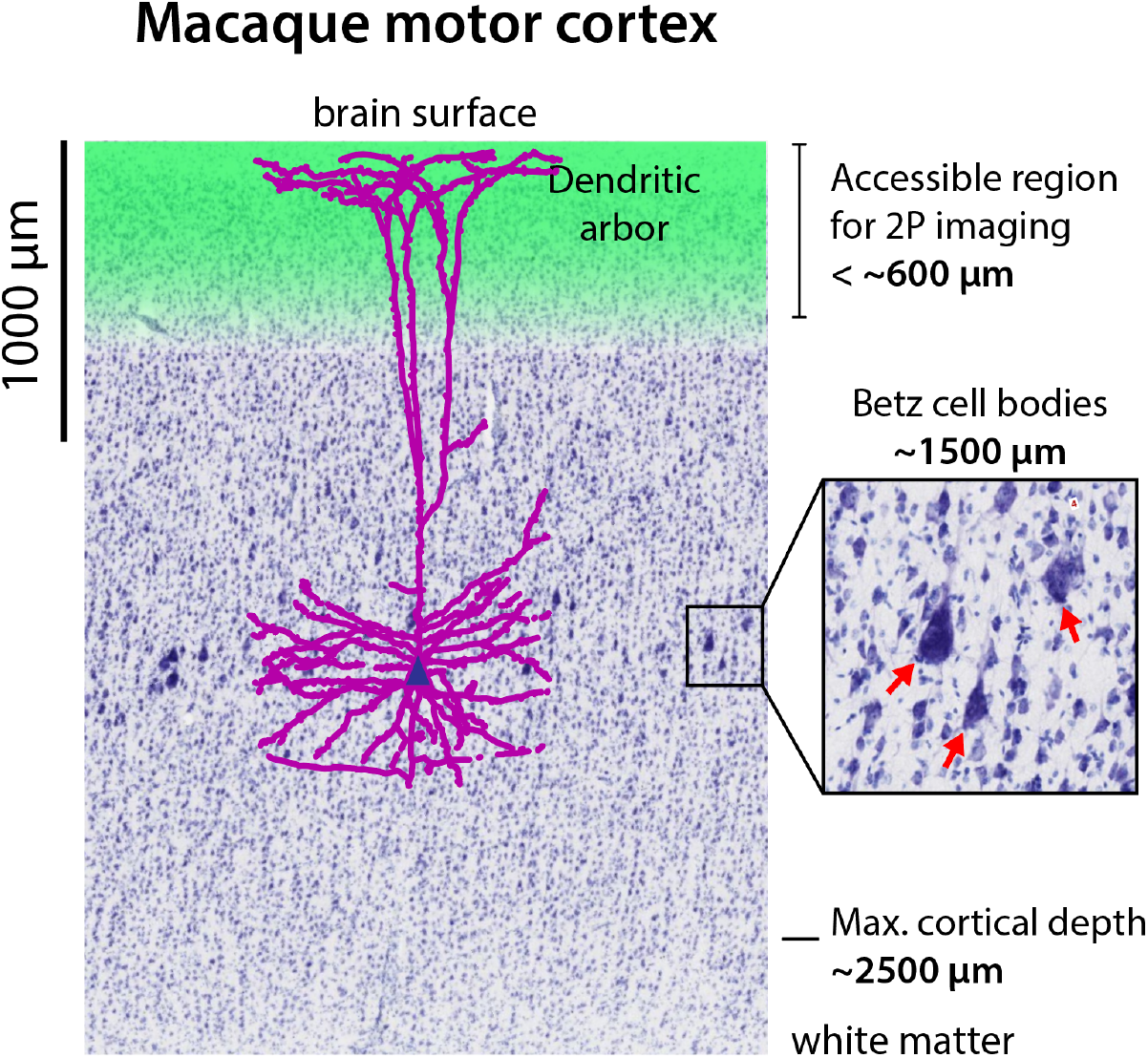
Dendritic calcium signals are readily accessible from layer V neurons. Two photon (2P) calcium imaging is currently capable of recording neural activity from the surface down to approximately 600µm, but photon scattering poses a challenge for imaging deeper. In this work, we demonstrate that it is possible to record neural activity of Layer 5 pyramidal neurons with cell bodies approximately 1500 µm below the surface (red arrows, inset) by imaging apical dendrites in superficial layers (purple), in addition to somatic signals from neurons located in layers 2/3 (blue cell bodies). Nissl stain source: brainmaps.org.

In this report, we sought to determine whether 2P calcium imaging at single-cell resolution of superficial neural processes in macaque motor cortex provides stable reaching-related signals sufficient to drive a real-time BCI. Achieving this experimental capability required combining multiple technical advances spanning implant design, optics, genetics, and low-latency computation (Fig. 2). First, we modified existing GCaMP6 constructs to promote cellular transport and promote expression of calcium indicators in the dendrites. Second, we co-optimized AAV serotype selection by immune-profiling individual macaques to improve viral transfection. Third, we designed a novel imaging implant to stabilize brain movement encountered when imaging in the motor cortex during monkey reaching behavior. We found that the forces produced during arm reaching were sufficient to induce significant tissue movement with conventional head restraint systems, presenting a greater stabilization challenge than faced in studies of macaque visual cortex^16–18^ or in marmosets^21, 22^. Coupled with the additional challenge of imaging fine neural processes in addition to somatic signals, our stabilization-optimized implant and head restraint system proved essential to imaging during motor behaviors.

**Figure 2.**
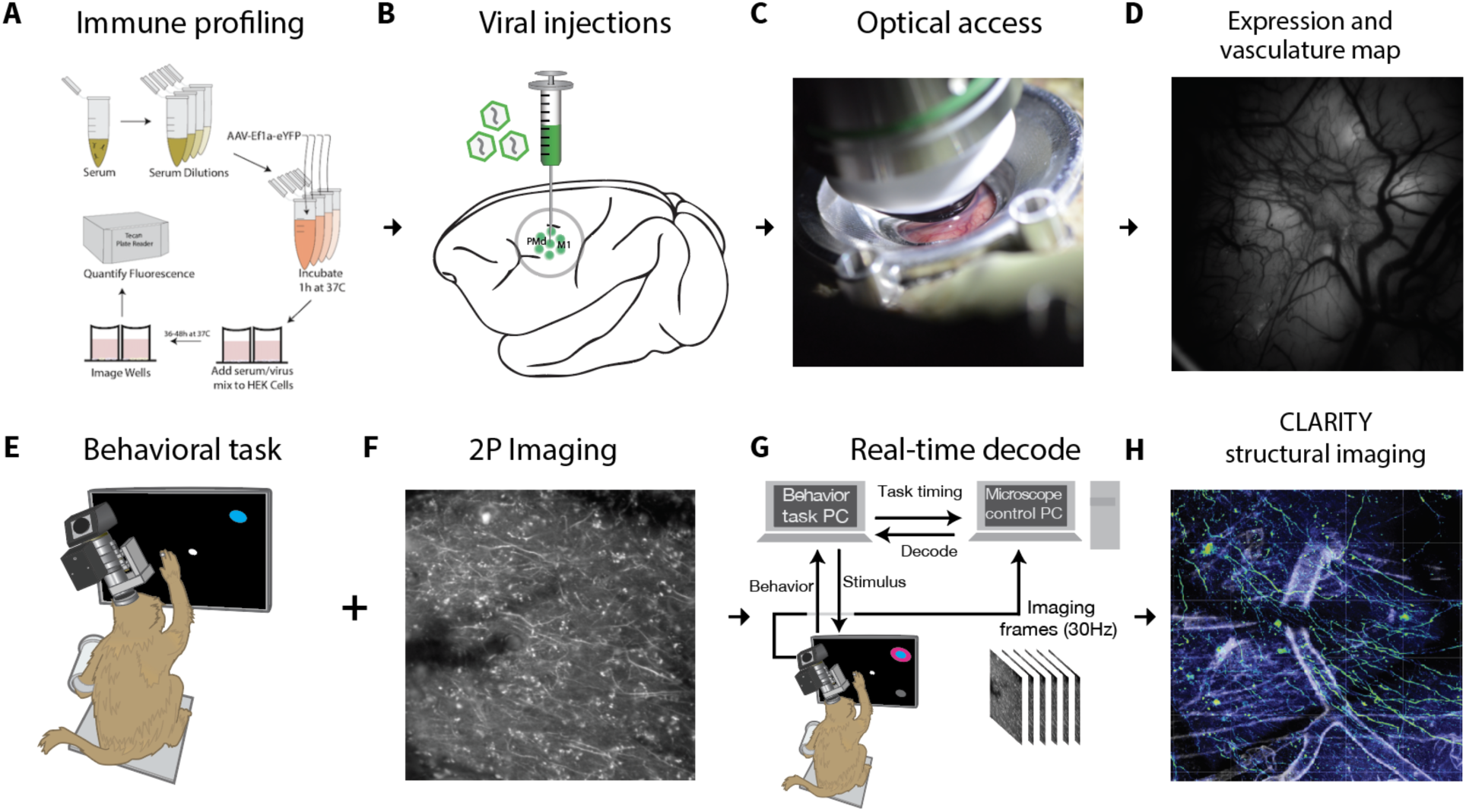
Experimental pipeline for combining functional imaging during motor behaviors with structural imaging in macaque monkeys. **(A)** Prior to imaging, we performed a neutralizing antibody assay in order to select an appropriate viral serotype tailored to the immune response of each monkey. **(B)** Viral constructs were injected into cortex to deliver the calcium reporter gene. **(C)** A chamber designed for chronic 2P imaging in premotor cortex and motor cortex was implanted. **(D)** Widefield (1P) imaging was used to assess GCaMP expression and establish vascular fiducial markers for navigating to specific sites on the cortex. **(E)** A macaque was trained to perform a reaching task to radially arranged targets. **(F)** 2P imaging was used to obtain functional signals at single-cell resolution from motor cortex. **(G)** During training trials, a decoder was trained on the imaging data obtained during reaching movements. Subsequently, during test trials, this decoder was run in real time to decode (predict) the reach target from 2P imaging data. **(H)** Post hoc CLARITY was performed to identify cell morphology, projection patterns and cell type.

These technical advances collectively enabled imaging of neural activity in macaque motor cortex with micron resolution across a large field of view, providing access to somatic and dendritic neural sources. We leveraged widefield (1P) imaging of vascular landmarks and GCaMP expression with 2P imaging to repeatedly localize the same neurons and dendrites across many experimental sessions and observed stable movement tuning in single dendrites over a timespan of weeks. We decoded these superficial neural signals in real time using an optimized, low-latency image processing pipeline to drive a real-time oBCI. We also localized functionally-imaged neural sources in a post mortem 3D CLARITY volume^34, 35^ and demonstrated that certain dendritic signal sources originated from layer 5 projection neurons.

This study serves as a proof of concept for combining emerging optical neural recording technologies with exploring a new neural compartment from which to gather BCI-relevant neural signals. This approach opens up opportunities for studying novel BCI technology designs^36^ (e.g., how best to design the placement of electrodes in a 3D volume to extract useful information, balancing tradeoffs between dense sampling of a small neural volume against broad sampling over a larger volume). It also demonstrates new capabilities for investigating how motor cortex controls arm movements by combining the benefits of optical imaging registered with post mortem anatomical imaging with awake, behaving NHP tasks, including the ability to make closed-loop experiment adjustments based on real-time neural read-outs^37, 38^.

## Results

### Obtaining optical access to motor cortex

Obtaining optical access to the brain in macaque monkeys is especially challenging for 2P calcium imaging due to the large size and short working distance of current multi-photon imaging objective lenses. 2P imaging requirements place constraints on the geometry of an implantable imaging chamber, while the implant must also provide tissue stabilization to minimize motion from pulse, respiration, and forces placed on the implant during natural behaviors in a motor task. Lastly, this chamber must remain sealed except during experiments but also provide easy access to the edge of the dura for cleaning and maintenance.

To address this, we developed an imaging implant that provided imaging access to a 12 mm diameter region of cortical tissue using a commercially available objective lens (Fig. 3A,B). This chamber used an implantable titanium cylinder and silicone artificial dura and was designed to balance the competing goals of maximizing the volume of imageable tissue while minimizing the craniotomy diameter and impact on the subject^39^. The chamber also provided simple access to the edge of the dura to remove new tissue growth and to allow for flushing fluid to reduce the risk of infection or to enable thorough cleaning if infections did arise^40^.

**Figure 3.**
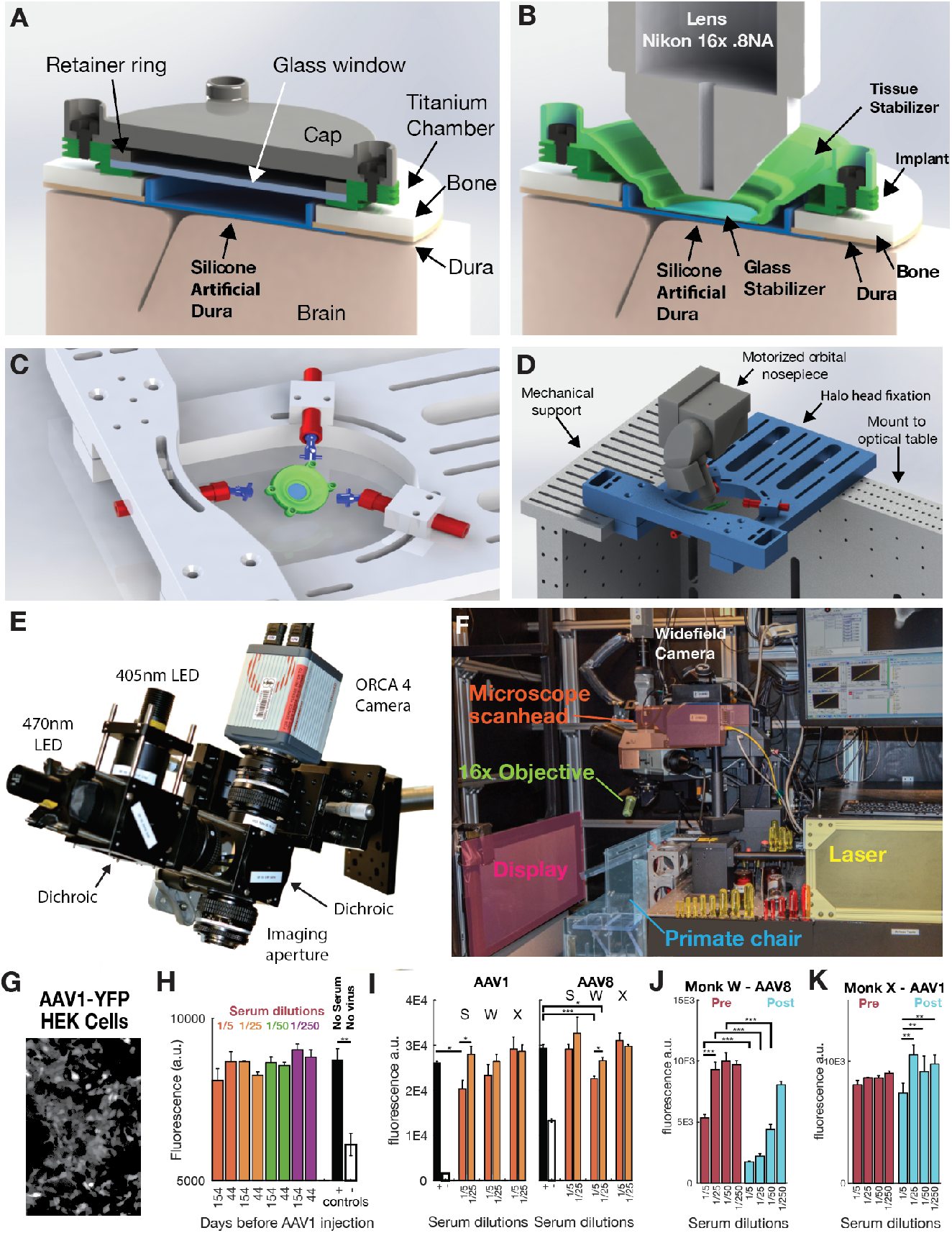
Implantable chamber, subject immunoreactivity profiling, and imaging apparatus. **(A)** Implantable titanium chamber in non-imaging configuration enabled observation through a glass window if the cap is removed. The glass window enabled long-term application of antibiotics and drugs to help maintain the health of the tissue margin. **(B)** During imaging, the cap and glass window were removed, and a temporary stabilizer is placed inside the chamber to restrict tissue motion via gentle downwards pressure on the surface of cortex. **(C,D)** While imaging, the implant was stabilized using three-point fixation to reduce motion of the tissue to micron levels. **(E)** Widefield imaging was performed using a custom microscope (see Methods) **(F)** During 2P imaging, a macaque sat in a standard primate chair in front of a stimulus display screen. The 2P imaging system is placed in a cantilevered position off the edge of the optical table to access the primate’s motor cortex. **(G)** Example image of AAV1-Ef1a-YFP infected HEK cells. **(H)** Neutralizing antibodies were stable across 100 days and multiple serum dilutions for monkey X (ANOVA, F_7,16_ = .96, p = 0.49) (n=3 replicates per experimental condition) **(I)** Each macaque subject (S, W and X) had pre-existing neutralizing to different AAV serotypes. **(J)** Monkey W developed significant antibodies to AAV8 post-injection (one-way ANOVA, F_7,16_ = 43.9, p < 0.0001). **(K)** Monkey X did not develop significant antibodies to AAV1 post-injection. (* p < 0.05, ** p < 0.01, *** p < 0.001). Mean + SEM represented in bar plots **H-K**.

Functional 2P imaging requires stabilizing neural tissue at the scale of microns during an imaging experiment. Changes in intracortical pressure due to pulse and respiration can cause periodic tissue motion inside the skull at the scale of many hundreds of microns, which presents a considerably greater challenge in macaques than in rodents and in smaller animals. In principle, fixed-depth rigid windows may be used to restrict tissue motion, as is comment in rodent experiments. In our experience, for chronic implants on the dorsal aspect of the skull, the brain may often recede from the widow over time rendering the stabilizing pressure of the window ineffective. To address this, we developed a removable tissue stabilizer, which mounts inside the recording chamber and uses a glass coverslip to apply gentle downward pressure on the surface of the artificial dura, restricting residual motion of the cortex (Fig. 3B).

Importantly, this pressure was only applied during imaging sessions, which reduced the potential for damage due to chronic focal compression. Design files for the imaging chamber and artificial dura components are provided in supplementary materials.

Micron-scale head restraint is crucial for successfully imaging at cellular resolution. Commercial primate head-fixation systems, designed for electrophysiology, are insufficiently stiff to prevent mechanical flexure at the scale of tens or hundreds of microns from the forces produced by natural arm movements (e.g., ∼1 kg arm, order of 1 m/s^2^ acceleration). We developed a highly rigid three-point head restraint system (Fig. 3C,D), which is conceptually similar to existing “halo” style head restraint systems^41–43^ but restricted head motion to just a few microns (Sup. Fig. S3). We integrated this stabilization system with an optical table capable of both widefield fluorescence imaging (Fig 3E) and 2P imaging (Fig 3F).

### Validating functional GCaMP expression and 2P imaging in motor cortex

Individual subjects will have different immunological states based on their particular exposure history to environmental viruses^44^. Although the brain has immune-privileged properties, whether pre-injection immunological status affects CNS expression of virally delivered constructs or elicits a systemic immune response is uncertain^45^. As such, viral infection and injection-related adaptive immune response may be highly variable between individuals, leading to consequences such as neutralization before infection, low expression of GCaMP, and/or deleterious systemic immune sensitization in certain subjects but not others. We employed a neutralizing antibody assay to quantify the levels of functionally neutralizing anti-AAV antibodies in serum samples. A series of blood serum concentrations (1/5, 1/25, 1/50, and 1/250) were incubated with AAV-YFP, and then added to wells containing HEK cells^46^, and YFP expression was measured with an automated plate reader. Decreased YFP expression indicated viral neutralization. Our goal was to select specific serotypes of AAV with the highest probability to infect neurons and the lowest probability of eliciting an immune response.

Before AAV injection, our assay could record stable pre-existing antibody levels across several months (one-way ANOVA, p = 0.49) (Fig. 3G,H). As expected, each macaque subject had a different pre-existing antibody status; monkey S was responsive to AAV1, monkey W was responsive to AAV8, and monkey X to neither (Fig. 3I). After viral injection, we found that monkey subjects with significant pre-injection anti-AAVs (e.g., monkey W) could develop immunoreactivity to small volumes of AAV viruses injected into cortex (Fig. 3J). In contrast, a low anti-AAV monkey (monkey X) demonstrated a response profile that seemed to mimic a primary exposure (Fig. 3K), consistent with a previous report^47^. The impact of immunoreactivity on the longevity of expression at healthy levels and on the success of subsequent additional injections remains to be determined.

Based on the neutralizing antibody assay results for monkey X, we identified AAV1 as a viral serotype with a high probability of success. We injected AAV1-CaMKIIɑ-GCaMP6f at two sites and injected two additional sites each with AAV1-CaMKIIɑ-NES-mGCaMP6f and AAV1-hSyn-GCaMP5G (Sup. Fig. S1). The decode results presented here were primarily imaged at injection sites for the AAV1-CaMKIIɑ-GCaMP6f construct. We monitored the expression of GCaMP after virus injection using wide-field and 2P imaging and observed first GCaMP expression at 4.5 weeks post injection (Fig. 4A, B). The AAV1-CaMKIIɑ-NES-GCaMP6f and AAV1-CaMKIIɑ-NES-GCaMP6f as well as primate-codon-optimized AAV1-CaMKIIɑ-mGCaMP6f and AAV1-CaMKIIɑ-NES-mGCaMP6f viruses were also tested in the visual cortex of monkey L, and functional signals were observed via widefield imaging in response to visual stimuli for all four viruses (AAV1-CaMKIIa-NES-GCaMP6f shown in Sup. Fig. S2, recorded using a separate imaging setup, see Methods and ^18^ for more details). Using vascular features identified via widefield imaging as fiducial landmarks, we located the sites of GCaMP expression with 2P imaging and observed neurons expressing GCaMP (Fig. 4E-G). These vascular features provided reliable landmarks allowing for imaging localization and reliable identification of individual neurons across multiple imaging sessions. These landmarks facilitated registration with *ex-vivo* CLARITY tissue-clearing (Fig. 4D and shown later in Fig. 6, 7).

**Figure 4.**
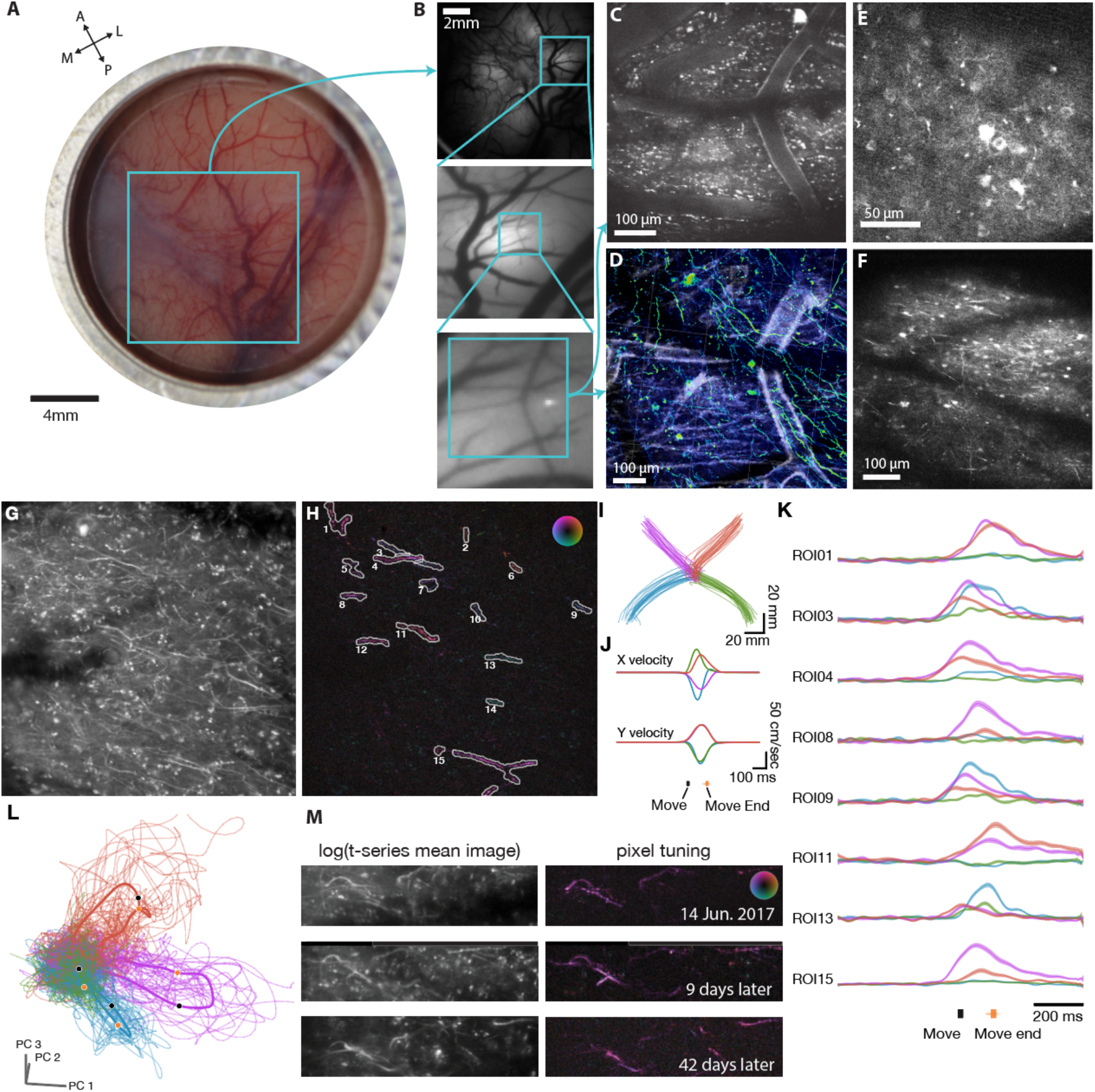
Multiscale, multi-modal imaging. **(A)** Imaging chamber with stabilizer in place under ambient illumination. **(B)** Cortical surface imaged using widefield (1P) imaging. **(C)** Region highlighted in cyan box in (**B**), imaged using two-photon imaging. **(D)** Same region as C, imaged ex-vivo using CLARITY and anti-GCaMP antibody labelling. **(E,F)** Two example fields of view including neural processes and L2/3 cell bodies ∼250µm below the surface vasculature (monkey W). **G)** Mean image over frames acquired during reaching behavior (monkey X, depth ∼100 µm). **(H)** Functionally defined regions of interest (ROI) identified from 2P time series imaging data, corresponding to neural features from (**G**). **(I)** Single-trial reach trajectories during center-out reaching with four radial targets. **(J)** Trial-averaged velocity profile in the X and Y axes for reaches shown in (**C**). **(K)** Trial-averaged neural responses ((ΔF/F) +/-SEM for a subset of high-SNR ROIs identified in (**B**). **(L)** Single trial neural trajectories, illustrating separation of trajectories based on target direction. Trajectories were calculated by projecting single-trial neural state into principal components 1-3. **(M)** The same neural processes observed during four-target reaching behavior is shown across three experimental sessions spanning 42 days.

**Figure 5.**
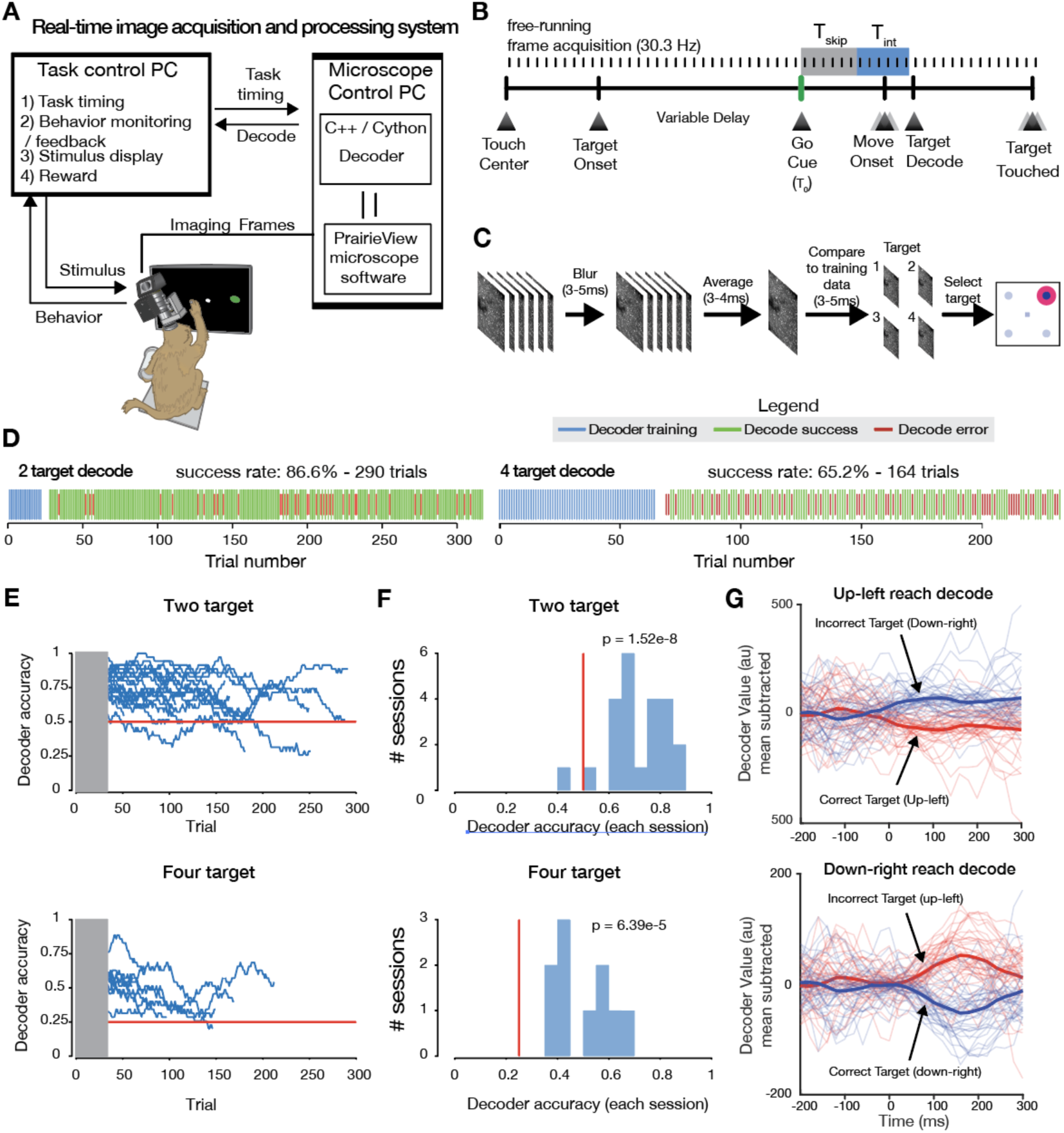
Real-time closed-loop decode of neural activity from functional imaging in motor cortex. **(A)** Real-time closed-loop stimulus control was implemented by decoding frames acquired from the microscope directly from memory buffers on the acquisition hardware. This provided rapid low-level access to imaging data to train and implement the decoder. Decode results were sent via ethernet UDP to the task and stimulus control computer. **(B)** Frames were acquired by the imaging system and are integrated by the decoder during a fixed window (T_int_) surrounding the movement onset beginning at a fixed latency (T_skip_) after the go cue. **(C)** The frames acquired during the integration time were blurred, averaged, and decoded using pixel-wise minimal mean squared error (MMSE) relative to training data (see Methods). **(D)** Timeline of oBCI decode sessions (monkey X). The decoder was trained using the first 20-40 trials in a given block (blue ticks). Subsequent trials were classified using MMSE decoder using raw pixel values as features. **(E)** Decode performance over the course of many individual sessions for 2 target (top) and 4 target (bottom) tasks (monkey X). Decoder performance was stable for up to hundreds of trials. In many cases, sessions were manually halted to record from a different field of view, not due to decreased decoder performance. Chance decoder performance is indicated by the dashed magenta line. **(F)** Histogram of mean success rate. Chance decoder performance is indicated by the magenta line. **(G)** Offline decoder value (cross-condition mean subtracted mean-squared error) using rolling 6-frame average for single trials.

**Figure 6.**
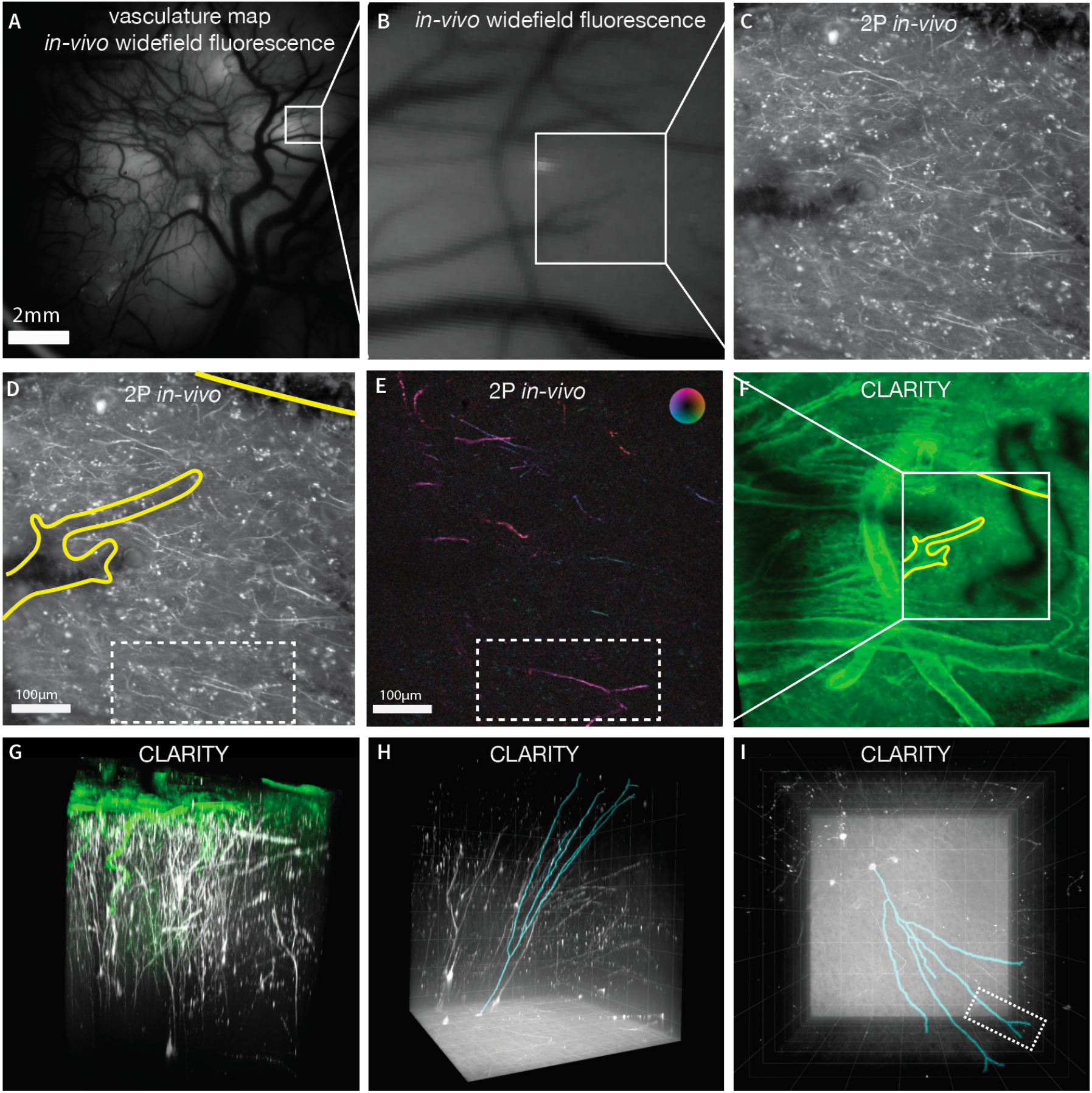
CLARITY registration with functional imaging. **(A)** A widefield fluorescence image revealing vasculature landmarks was used to locate and register 2P imaging FOVs. **(B)** Higher-magnification FOV from (A). **(C)** Example mean intensity projection of all 2P frames acquired during a decode session using the FOV indicated in (B) by blue box. **(D)** The same image as (C) with identifiable vasculature features used for FOV alignment highlighted in yellow. Selected dendritic process are outlined in white dashed rectangle. **(E)** Pixel-wise tuning map, same FOV and presentation style as in Figure 4H. **(F)** CLARITY volume showing wide-area surface vasculature (lectin stain, green) with FOV from (C-E) highlighted in yellow. The dark features on the right of the image is an imaging artifact from an occluding object only present in the CLARITY imaging. **(G)** Rendering of CLARITY volume showing a close up of neural processes extending from the FOV in (A,B) down to areas below the imageable regions using *in-vivo* 2P imaging (green: lectin, white: GCaMP stain). **(H)** Rendering of CLARITY volumetric imaging showing a side view of motor cortical tissue spanning cortical lamina with putative Betz cells located in Layer 5, approximately 1500µm below the surface. Blue outline indicates the traced reconstruction of the dendritic process imaged superficially in (E), traced from the dendritic arbor down to the cell soma. The location and large size of the soma suggests this cell is likely to be a Betz cell. **(I)** Surface image of reconstructed arbor, white dashed rectangle indicates matching region from (E).

**Figure 7.**
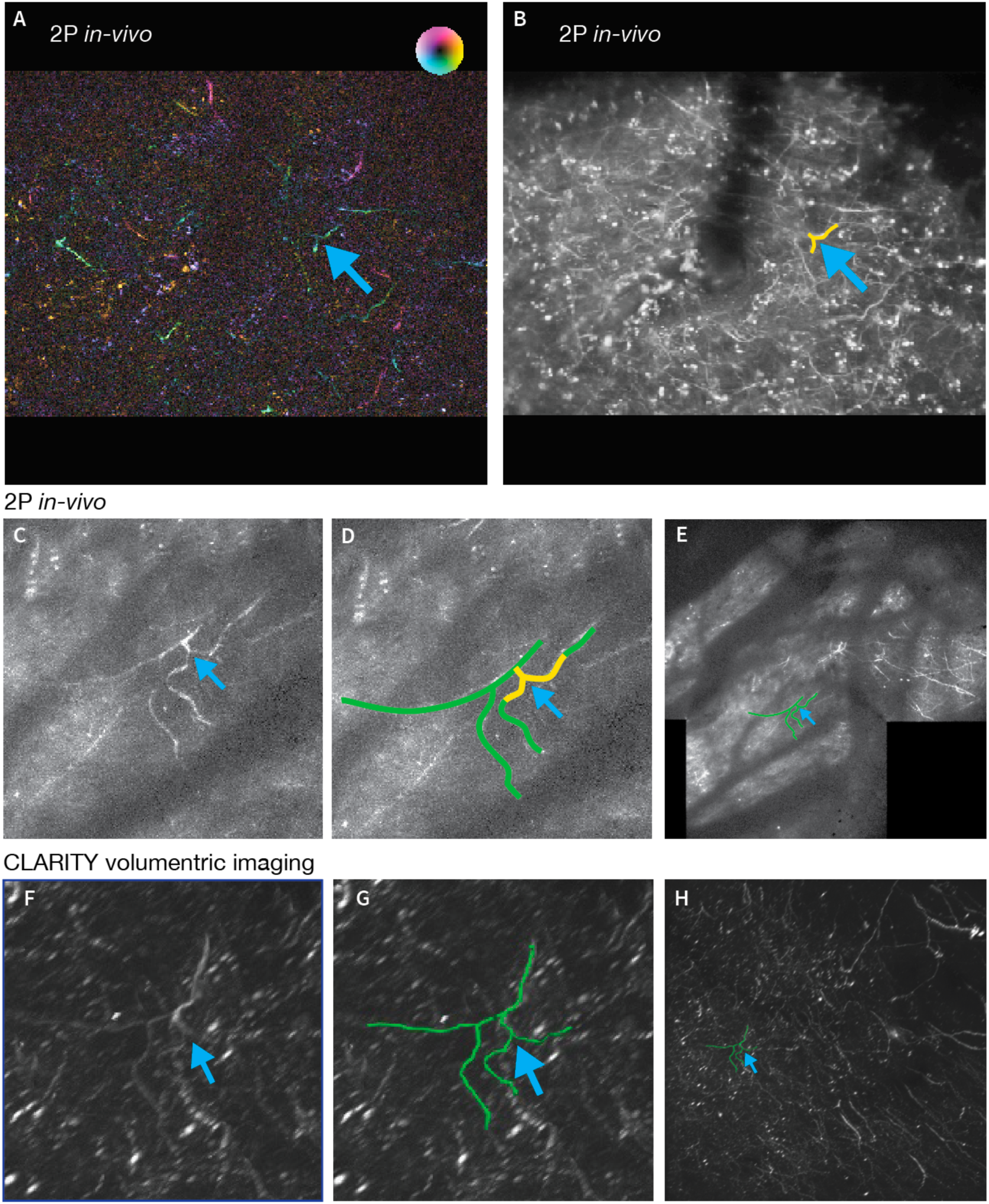
A second example a neural process functionally imaged and reconstructed in the registered CLARITY volume. **(A)** Pixel-wise tuning map, same presentation style as Figure 4H. A neural feature of interest is indicated by the blue arrow **(B)** Mean intensity projection of FOV in (A). The same process is labelled with a blue arrow. This image represents only a thin slice through the tissue volume. **(C)** Maximum intensity projection from *in-vivo* volumetric z-stack showing projection of the neural features labelled in (A) and (B). More structure is present in this image than in (B), since this image is a maximum intensity projection of images acquired at multiple depths **(D)** Same image as C, with the neural feature of interest from (A-C) traced in green. **(E)** Maximum intensity projection of stitched 3D volume assembled from *in vivo* 2P imaging. **(F)** Closeup view of *post-hoc* imaged CLARITY volume. The neural feature from (A-E) is marked with the blue arrow. **(G)** Same as F with the feature from (A-E) traced in green. **(H)** Wide view of CLARITY volume.

2P imaging fields of view included all fluorescent signals present in superficial cortical layers, down to approximately 550 µm, including somatic signals in superficial cortical layers (Fig. 4E, F) as well as calcium signals from GCaMP-expressing dendrites (Fig. 4G). In several injection sites across two subjects, we also observed bright fluorescence from some cell bodies which appeared to have filled nuclei and non-modulated signals from somas located between 0-550 µm from the surface of cortex and appeared within five weeks of injection (Sup. Fig. S5).

Signal contributions from individual ROIs were identified using pixelwise correlation with hand velocity, watershed segmentation, followed by manual inspection (Fig. 4H). Functional modulation of GCaMP expressing cells in the caudal aspect of the dorsal premotor cortex (PMd) and adjacent gyral M1 during motor behaviors was assessed using an instructed-delay reach task, which elicited rapid, straight reaches to each of four radially positioned targets (Fig. 4I, J). Arm movement was preceded by strong modulation of the calcium signal in neural processes. We constructed a pixelwise tuning map by regressing functional ΔF/F responses against hand velocity (Fig. 4H); individual neural processes exhibited consistent tuning for a specific reach direction within each ROI (Fig. 4K). Trial-averaged neural responses generally exhibited increases in activity preceding hand movement, consistent with the so-called condition (reach direction) independent signal widely reported in motor cortex^48^. Applying principal components analysis (PCA) to single trial neural responses from all ROIs yielded trajectories in neural-population state space^14, 49^ which traveled along distinct paths for reaches to the four different targets (Fig. 4L), consistent with the tuning observed in individual ROIs. This is also consistent with what we and others have reported previously using action potential emission rates measured from motor cortex with electrode arrays^50^. We returned to the same fields of view to image the same neural processes multiple times over several experimental sessions, spanning up to 42 days, and observed stable reach tuning in all neural processes for which we obtained data from multiple sessions (Fig. 4M; Sup. Fig. S4).

### Real-time decoding of reaching from 2P imaging of dendrites

To demonstrate real-time decoding of arm-movement behavior from functional neural imaging responses, we implemented a real-time, discrete target decoder as has been previously developed using electrophysiological recordings^7, 8, 51^. Here the discrete-target oBMI is driven by a low-latency image processing pipeline, trained to decode reach direction dendritic calcium signals (Fig. 5A). The majority of GCaMP-expressing neurons with functional signals in this experiment were located in gyral M1, caudal to premotor cortex. As anticipated, for activity in M1, as opposed to in PMd, we did not observe sufficient preparatory activity in this region to facilitate decoding the upcoming reach target from delay period signals^7, 8^.

At the start of each decoding session, training data were obtained by storing 2P imaging frames during the peri-movement period of natural arm reaching for 8-15 trials per target direction (Fig. 5B). We used the trial-averaged peri-movement imaging frames from the training data to train the decoder (Fig. 5C). After this training period, target classification was performed by analyzing frames during the same peri-movement integration window (T_int_), which began at a fixed delay after the “go cue” (T_skip_) instructing the monkey to reach. We compared these frames with the per-reach-direction template frames learned during training using a minimum mean squared error decoder (see Methods). The decoded target was subsequently sent to the task controller and a visual cue was presented to the monkey to indicate decode success or failure. A liquid reward was also provided after successful decode trials. The total processing time after frame acquisition was less than 15 ms (<1/2 of the imaging-frame period, measured from end of final galvanometer scan line), and the decoded target was visually rendered on the screen prior to the monkey touching the target with his hand. We note that in this experiment, the decoded target presentation did not (at least directly) affect the monkey’s motor output, but instead served as a demonstration of a functioning, real-time optical behavioral decoder where the signal-to-noise ratio of the optically-recorded neural population was sufficiently high even with a brief integration period (T_int_).

We assessed the online decode performance for selecting the correct target, from either two or four possible targets, and were able to achieve up to 86.6% successful decode for two targets (p = 2.17e-39, two-sided binomial test, chance level 50%, 290 trials) and 69.9% successful decode for the four-target case (p = 9.99e-16, two-sided binomial test, chance level 25%, 73 trials) (Fig. 5D). Decoder accuracy for single sessions remained above chance for hundreds of trials (Fig. 5E), and aggregated decoder performance for all sessions was significantly above chance (p = 1.52e-8 for the 2 target task, 23 sessions across six days, rank sum test; p = 6.39e-5 for the 4 target task, ten sessions across six days, rank sum test; Fig. 5F). Decoder confidence could also be assessed over time by assessing the difference in decoder score for frames throughout single trials. These signals diverged around the time of movement onset and remained separated through the duration of movement (Fig. 5G).

### Registering functionally imaged cells with anatomy via CLARITY

Having demonstrated that dendritic calcium signals are modulated during movement and are capable of driving an oBCI, we sought to identify the somatic sources of these signals. Using fine-scale brain vasculature as fiducial markers, it was possible to register and align imaging data from multiple fields of view and across multiple imaging modalities to within a few tens of microns within the tissue volume (Fig. 6A-D). These vascular features provided consistent reference points for navigating to and recording from the same population of cells across multiple sessions, as shown in Fig. 5. These same vascular features also enabled precision alignment between *in vivo* 2P calcium imaging and *ex vivo* volumetric imaging, enabled by clearing a large tissue volume containing the entirety of motor cortex using CLARITY. Within this CLARITY volume, we verified that many GCaMP-expressing deep layer 5 cells extended apical dendrites towards the cortical surface. We then traced the detailed morphology of one particular process imaged during an oBCI decoding session (white dashed rectangle, Fig. 6D, E, I). This tuned process was a branch of an apical dendrite of a cell originating approximately 1500 μm below the surface of cortex (Fig. 6F-I). The cell body was approximately 60 μm in diameter, suggesting that this cell is likely to be a Betz cell, a class of large upper motor neuron which projects to the spinal cord^52^. We demonstrate a second example of registering features across imaging modalities in Fig. 7, in which clearly identifiable features are identified in 2P *in vivo* functional imaging, 2P *in vivo* volumetric z-stacks, and *post-hoc* 2P CLARITY imaging.

## Discussion

We developed an all-optical motor BCI driven by calcium signals, enabled by a suite of engineered optimizations for stable 2P imaging in rhesus macaques engaged in motor tasks. The optical window, as part of the custom-designed implant, affords imaging access to many hundreds of thousands of neurons across PMd and M1 and is compatible with other brain regions. Using this implant, we achieved stable, chronic 2P imaging of motor cortical neurons in awake, behaving macaques. We demonstrated that signals imaged from dendrites located in superficial layers were modulated during movement, were directionally tuned, and exhibited sufficient signal-to-noise ratio to enable a real-time oBCI using closed-loop image processing to decode a monkey’s behavior from neural activity to provide low-latency visual feedback and reward. We leveraged wide-field imaging of vascular fiducial markers to return to the same neurons and observe consistent direction tuning in neuronal processes over multiple weeks. Lastly, we co-registered processes imaged in 2P with post-mortem CLARITY volume images, enabling the tracing of individual processes back to their neuronal soma, thereby demonstrating that some of the imaged dendrites likely originated from layer 5 corticospinal projection neurons.

Importantly, we do not suggest that imaging is necessarily a viable recording modality for clinical BCIs in the near term due to the need to introduce exogenous calcium reporters to the human brain using viral vectors. Instead, we view optical imaging and oBCIs as an important pre-clinical tool to address critical questions that are challenging to address with electrical recording alone^36, 53^. Optical imaging and closed-loop oBCIs complement electrophysiology methods by addressing several limitations of current array technologies, such as providing the potential to accurately track large neural populations over timescales longer than several days, densely recording from 3D volumes of tissue, and merging genetic neural circuit dissection techniques with BCI designs to better understand the neural substrate for neural prosthetic control. Even high-density silicon electrodes (e.g., Neuropixels probes^25, 54^) provide limited volumetric tissue coverage, cannot easily be moved once inserted, and lack cell-type information and ability to target genetically-specified cell types. Moreover, the process of engineering appropriate electrical stimuli for sensory write in^55–57^ would greatly benefit from direct optical observation of the complex stimulation-evoked patterns of neural activity^58, 59^. Ultimately, imaging methods will deepen our understanding of the basic science of natural movement control and will help advance BCI algorithms, enabling the development of next generation BCIs for animal research and human clinical use.

Achieving stable 2P imaging in monkeys engaged in motor behaviors necessitated several technical advances. First, we developed an optical window optimized for large multiphoton lenses and an accompanying stabilization system to obtain optical access to cortex, which is more challenging in macaques than in rodents or smaller organisms. Stabilization proved more challenging in frontal cortex than in the occipital lobe (e.g., ^17, 18^) due to the effect of gravity, which pulls the brain away from the dorsal imaging plane. The artificial dura system facilitated reliable optical access for many months by maintaining an environmental seal and enabling frequent cleaning to minimize infection risks for chronic implants after removing the dura. Once optical access had been established, the adjustable tissue stabilization and three-point head fixation system enabled stable imaging despite substantial cardiac and respiratory-driven brain pulsation as well as brain-in-skull motion caused by the monkey’s movements. Additionally, the artificial dura window designed provided access to an 18 mm diameter region of cortex, which was considerably larger than the field of view that our 2P imaging system could simultaneously image. Our implant design could also be used with wide field of view imaging technologies such as the mesoscope to address an expanded field of view, thereby vastly increasing the number of simultaneously recorded neurons^23^.

Second, we obtained functional expression of GCaMP6f throughout motor cortical neurons including apical dendrites. The robust, functional expression of GCaMP in dendrites may have been aided by designing a macaque codon-optimized GCaMP which included a nuclear export signal (NES) target peptide^60^. The probability of achieving successful expression might also have been improved by performing a neutralizing antibody assay to pre-screen viral serotypes against subject-specific immune status. Serotype, selection of genetic promoters, and the viral delivery protocol are all likely to play an important role in determining whether expression levels are healthy and effective. We found that small intracranial injections of AAV could elicit a humoral immune response in pre-sensitized subjects, as measured by the neutralizing antibody assay. While this information can guide the selection of candidate viral serotypes to minimize anti-AAV immune response, more work is needed to determine whether humoral responses to AAVs negatively impact neuronal GCaMP expression. We demonstrated functional expression of GCaMP using AAV1 in macaque motor cortex and validated safe and effective expression of these constructs in a second subject in V1.

Third, we engineered a low-latency image processing pipeline to train and execute the real-time optical decoder which provided the movement estimates needed for the oBCI system. This pipeline leveraged memory buffers exposed by the microscope hardware and image processing code optimized to achieve low latency to achieve an online decode, which is conceptually similar to a 1D imaging decoder reported in mice^61^. This enables a large class of new experiments in which neural activity is read out and used to modify the task or stimulus in real time, e.g. using a BCI decoder to study motor adaptation to visuomotor rotation^62^. Combining this capability with genetic targeting can elucidate the roles of different cell types in contributing to motor adaptation and control.

Finally, to identify the source of a subset of the dendritic signals we imaged, we optimized CLARITY to clear and immunostain a large volume of the macaque brain, encompassing the whole of motor and premotor cortices. By localizing blood vessels in both the cleared volume and functional datasets, we were able to co-register between functional datasets acquired with 2P and volumetric CLARITY data, exploiting neuronal morphology visualized through GCaMP expression in dendritic arbors to match specific cells. We used this technique to validate the layer 5 somatic source of specific dendritic processes which drove the oBCI decoder.

### Identifying cell types of recorded neurons

Electrophysiological recordings are limited in their ability to connect descriptions of neural population dynamics to the underlying neural circuit structure. In particular, using electrophysiology alone, it is not currently possible to study how coordinated population activity arises from the constituent neuronal cell types, the fine spatial organization of cells within the circuit, and connectivity of the network^14, 63, 64^. Several studies have used spike waveform shape to distinguish putative excitatory and inhibitory neurons^65–67^ and with increasing electrode density finer details of each neuron’s electrical image onto many nearby channels can better resolve additional cell classes^68, 69^. However, it is difficult to validate the accuracy of this approach. Genetic approaches can bridge this gap and provide richer information on the diversity of neurons present throughout cortex^70–72^.

In our datasets, we localized and traced several functionally-tuned cells within the CLARITY volume with high confidence in the correspondence between *in vivo* and CLARITY imaging. One cell could be identified as a putative Betz cell due to its unique morphology. Once a broader panel of labeling methods are validated in rhesus macaques and tissue clearing techniques are better optimized for macaque and human tissue^73^, we anticipate that the cell-type information provided by co-registered CLARITY volumes will rapidly increase. By providing access to both structure and function, optical methods can discover connections between computational-level descriptions afforded by a dynamical systems view of neural population activity and the underlying mechanisms that generate and shape these dynamics to drive behavior. In motor cortex, this opens the door for further work investigating the role of distinct cell types in motor control, an approach which has been highly successful in the mouse^74, 75^.

### Imaging Neural Projections

The majority of previous calcium imaging experiments used isolated neuronal somas as the primary signal source. However, in rodents and larger mammals, imaging neuron cell bodies below superficial layers of cortex (down to ∼600 μm below the surface) at standard frame rates (∼30 Hz) is not currently possible using standard 2P imaging technologies due to photon scattering and absorption. It is not currently possible to image somatic calcium signals from layer 5 cells in macaque motor cortex, which may be located as deep as 1.5-2.5 mm from the surface, without using a penetrating GRIN lens^76^ that causes damage to the adjacent neural tissue^27^. The approach we demonstrated here accessed calcium transients from these deeper neurons through their apical dendrites, which arborize near the cortical surface. The data we presented demonstrate that superficial processes in M1 exhibit directionally-tuned movement modulation, consistent with signals from extracellularly-recorded motor cortical action potentials^25^. This correspondence supports additional work to leverage these dendritic calcium signals to explore neural population activity during motor behaviors. Moreover, this approach could also be used to study computations in active dendrites with calcium reporters^31, 32^ or with fast, genetically encoded voltage indicators^77^.

More broadly, imaging of superficially projecting axons could be used to study a variety of neuromodulatory influences on cortical processing. Previous studies in rodents have shown that it is possible to optically record activity from GCaMP-expressing neuronal projections^78, 79^. This suggests that by expressing GCaMP in subcortical areas, which source a specific neuromodulatory influence on cortex, the activity of these axonal projections may be imaged directly at their cortical target. For example, dopaminergic afferents from ventral midbrain laminate throughout cortex including superficially in layer 1^80^. Imaging of cholinergic afferents from basal forebrain, known to activate layer 1 inhibitory circuits^81^, could elucidate the mechanisms of signal enhancement in primary visual cortex^82, 83^ and in memory function in prefrontal cortex^84^. Analogous approaches could be used to study the influence of GABAergic inputs from basal ganglia into frontal cortex on reward learning and coordination of motor actions^85^ as well as thalamic afferents^86^ and cortico-cortical connections^87^ with superficial synaptic targets.

### Stable tracking of neural populations across sessions

In macaques and other NHPs such as marmosets, there is a growing interest in tracking neural populations over time to address a collection of learning-related questions. Multielectrode array recordings in M1 and PMd have been used to study population changes in motor cortex during visuomotor adaptation^62^ and force field adaptation^88^. Researchers have employed carefully designed perturbations to BCI decoders—in which the behaviorally-relevant readout axes of neural activity are controlled directly by the experimenter—to probe constraints on learning imposed by neural circuitry^37, 89, 90^. Within the context of BCI, researchers are also developing ‘co-adaptive’ BCI algorithms, in which a subject learns to improve BCI performance by generating specific patterns of neural activity via neural plasticity, while the decode algorithm concomitantly adjusts to make the BCI easier to control (e.g., ^4, 91^; reviewed by^3^). Understanding and facilitating co-adaptation is generally believed to be an important next step in BCI design^92^.

Multielectrode arrays, however, are susceptible to waveform drift caused by micron-scale shifts in electrode placement. Tracking the activity of a specific population of neurons across multiple experimental sessions spanning days or weeks is quite challenging^93–97^. Precise tracking of large populations of neurons using imaging could be leveraged to study learning-related changes at single-cell resolution across longer timescales. Optical techniques can meet this need by recording from the same population of neurons across sessions. This is possible because morphological features of the neurons allow for high confidence in identifying the same neurons in multiple images. Using a combination of widefield and 2P imaging, we were able to localize the same field of view using vasculature and GCaMP expression patterns as fiducials, which enabled recording the same neurons in multiple sessions. This capability underscores the utility of functional imaging for studying the evolution of neural population dynamics over extended timeframes, particularly for experiments involving across-session learning or long timescale adaptation. For example, 2P imaging has proven particularly useful in rodents for observing learning-related changes in neural population activity^98–103^, and we believe a similar approach may be employed in primates.

### Conclusions

The technical advances reported here enabled stable imaging sufficient to record direction-tuned signals in a population of dendrites and to drive a real-time direction decoder. The observed accuracy rates for the online oBCI decoder were significantly above chance but are not yet at parity with results obtained using multielectrode electrophysiology methods (e.g., Utah arrays^8^). While our proof of concept oBCI was a discrete decoder for reaching direction, previous simulations have shown that it is possible to decode continuous hand movement trajectories from simulated optical signal, despite calcium indicators having (1) relatively slow response kinetics to action potentials, (2) various sources of noise, and (3) non-linear relationship, including saturation, between the neural spikes and fluorescent signal^104^. Additionally, our analysis in that study indicated that oBCI decoding performance should significantly increase with improvements in the temporal resolution of calcium reporters^105^ or with high-SNR, genetically-encoded voltage indicators^77^.

The GCaMP expression we observed was sparser than comparable results using virally-transfected GCaMP in mouse^106–108^, marmoset^20, 22^, and primate V1^17, 109^ for most injection sites. Future applications would benefit considerably from denser and more reliable expression of the reporter construct.

Fortunately, considerable advances in macaque optogenetics have provided insight into effective delivery vectors for achieving expression^110^. Reliably obtaining and maintaining healthy levels of GCaMP expression in motor and premotor cortex remains challenging and is particularly important in highly trained macaques. Future experiments may benefit from subject-specific serotype pre-screening as performed here, careful consideration of serotype and promoter^111^, and precision titration of expression (e.g., via tetracycline-gating) to maintain robust, functional signal^20^. We anticipate that achieving denser expression will enable more sophisticated decoding approaches capable of driving the continuous velocity of a cursor or a robotic arm.

This work demonstrates new capabilities for combining emerging neural recording technologies with closed-loop BMI and *post-hoc* volumetric, structural imaging. This pipeline provides new opportunities to leverage genetic neural circuit dissection techniques within motor neurophysiology studies and brain machine interface experiments. Such an integrative approach is needed to elucidate the neural circuits that control natural movements and neural prosthetic devices.

## Methods

### Animal Subjects

All procedures and experiments were approved for animals S, W and X by the Stanford University Institutional Animal Care and Use Committee (IACUC), and for animal L by the University of Texas at Austin IACUC and were performed in compliance with the Guide for the Care and Use of Laboratory Animals. Three male rhesus macaques (X, S and W) were used for two-photon imaging experiments and a fourth rhesus macaque (L) was used for secondary validation of virus constructs in V1.

### Surgical procedures

In monkey X, we implanted the imaging chamber in a sequence of two surgeries in order to (1) minimize the duration of surgical procedures, and (2) allow for behavioral training with head fixation prior to opening the dura. After the dura was opened, the surface of the cortex required intermittent cleaning and maintenance, and an opaque “neomembrane” often began to grow over the surface of the cortex approximately 2-4 months after opening the dura. In the first surgery, we implanted the chamber over the surface of the bone, sealing the chamber to bone interface with C&B Metabond dental cement (Parkell), and cementing the chamber in place using Palacose bone cement (Heraeus Medical) and titanium bone screws (Synthes Inc.). Custom-machined headposts were implanted to allow for head fixation during behavioral training. Several months later, after completing behavioral training, we performed a second surgery to remove the skull (in the region of the cylinder) and dura from the center of the 2P chamber, to perform viral injections and to place the artificial dura (AD). Viral constructs were injected using pulled glass micropipettes (∼25 μm tips), beveled using a micropipette grinder (Narishige EG-401 pipette beveller) using a nanoliter injector (WPI Inc.). To assist in visualizing viral injections, trypan blue dye (0.4% (w/v)) was diluted 1:10 in saline and mixed with the viral suspension^18^.

For monkeys S and W, we implanted an earlier generation of titanium chamber design, which threads into a craniotomy. For these surgeries, the imaging chamber was implanted in a first surgery, but the dura was left intact, and virus injections were performed using stainless steel syringes (Hamilton) outside the operating room while the monkey was performing a behavioral task. After waiting for expression, the dura was then resected, and the AD placed in a second surgery. While this approach has the advantage of not exposing the surface of the brain while waiting for GCaMP expression (∼8 weeks) prior to imaging, it required injections to be performed through the dura, preventing careful targeting of virus as the vasculature on the surface of cortex was not visible. This led to additional uncertainty with regards to injection location and depth, made it impossible to avoid surface vasculature while injecting, and increased the required volume of virus injected in order to be confident that a sufficient volume of virus was injected in the target lamina. As such, in monkey X we adopted the more targeted approach, performing virus injections using glass micropipettes with the surface of the brain exposed, which is the preferred approach for new experiments^18^.

### Implant design and maintenance

The imaging chamber design strikes a balance between (1) enabling imaging access to a large volume of tissue (at least 12 mm in diameter up to ∼ 1000 um deep) using commercially available multiphoton objective lenses and (2) minimizing the implant size. The implant must allow for stable head fixation during reaching behaviors and allow for simple replacement of the silicone AD and access to the edge of the dura for routine cleaning and maintenance. These design constraints suggest a large-diameter but low-profile imaging chamber and multi-point head fixation. Design files for the chamber and associated hardware are provided in supplemental materials.

### Head restraint and implant immobilization

Initial testing revealed that single-point head fixation methods, as are commonly used for electrophysiology, were incapable of restricting tissue motion at the scale of microns. We tested a three-point acrylic-free footed headpost system that uses bone screws to secure headposts to the skull and found that this too was not sufficient to restrict micron-scale implant motion during arm-movement behavior. Instead, we found that using a single larger implant constructed with bone cement (Palacose, Zimmer BIOMET Inc.) and titanium mandible straps (Synthes Inc.) fixed to the skull with bone screws allowed for more rigid fixation by distributing the loads between the multiple fixation points through the bone cement. Design files for the head restraint and associated hardware are provided in supplemental materials and details provided upon reasonable request.

### Tissue stabilization

Prior to imaging, the outer window and retaining ring were removed under sterile conditions, and the AD was exposed to the air. The tissue stabilizer was placed within the chamber, placing gentle mechanical pressure on the top surface of the AD. The tissue stabilizer consisted of a conical aluminum component which sat inside the chamber and held a circular glass coverslip against the artificial dura. Since the location of the surface of the brain could change over time due to tissue growth, recession, or other factors, we fabricated a set of tissue stabilizers at different fixed depths in 500 µm increments. Sterile saline was placed in the chamber prior to placing the stabilizer as an index matching fluid between the silicone and stabilizer glass.

### Maintenance and tissue cleaning

After an experiment, the tissue stabilizer was removed, and a solution of agarose and vancomycin was applied to the dura edge, typically every one to four days. Physiosol (Pfizer Inc.) with added vancomycin was applied to fill the remaining chamber volume, and the glass window was secured in place with the retainer ring, and the chamber cap is secured on top. Under normal conditions, the AD did not need to be regularly removed or replaced for cleaning. Over timescales of several months, we observed ‘neomembrane’ tissue growth under the AD but over the surface of the cortex. Over time, this tissue grew in thickness, blocking optical access to the cortex, and requiring careful surgical dissection and removal. In practice, the timeline for tissue removal could be variable but was usually required every 2-4 months to retain imaging performance. The wound margin at the edge of the durotomy also experienced tissue growth and required periodic trimming to prevent excessive buildup of tissue above the artificial dura.

### Widefield Imaging in V1

As part of the development and verification process for the GCaMP constructs, we validated some constructs using widefield imaging in visual cortical areas in monkey L (Sup. Fig. S2). We used a large (6 x 6 deg^2^) sine wave grating at 100% contrast with a spatial frequency of 2 cpd. The mean luminance of the screen was set at 30 cd/m^2^. The grating was flashed with a temporal frequency of 4 Hz, (100 ms on, 150 ms off) while the monkey was performing a fixation task. The behavioral task and widefield (1P) GCaMP data analysis in the rhesus macaque (monkey L) were performed as described previously^18^.

### Two-photon imaging in premotor (PMd) and primary motor (M1) cortex

Imaging was performed using a Bruker Ultima *in-vivo* microscope with a custom motorized orbital nosepiece (Bruker Inc.) to provide off-axis imaging with a Nikon 16x 0.8 NA objective lens. Images were acquired at 512 x 512 resolution (or occasionally at lower resolution while imaging a portion of the 512 x 512 pixel field) at a single depth at 30.3Hz using resonant-scanning galvanometers. Behavioral sessions lasted between 90-180 minutes and no bleaching of GCaMP6f was observed over this time.

### Behavioral task and decoder

Monkey X was trained to make point-to-point reaches with the arm contralateral to imaging implant on a delayed center-out-and-back task in a vertically-oriented 2D plane as described previously ^112^. The monkey initiated each trial by holding a point at the center of a display screen, placed approximately 30 cm from the eyes. Next, one of 4 radially-arranged targets appeared 10 cm from the center and jittered during the delay period (randomized period, ranging from 100-700 ms). Next the target ceased to jitter, and the central fixation point disappeared, thereby indicating a ‘go cue’. The monkey was free to initiate a reach following the ‘go cue’, with the time between the ’go cue’ and the detection of movement termed the reaction time (RT). To exclude rare trials where the monkey may have anticipated the ‘go cue’ (i.e., RTs < 180 ms) or where the monkey may have been distracted (i.e., RTs > 620 ms) we aborted trials in real time by blanking the screen and withholding reqard if the RT was outside of the 180-620 ms RT range. Hand position was measured in 3D and in real time with a Polaris infrared bead tracker (Polaris, Northern Digital, Ontario, Canada) which samples 60 times / sec and has < 1 mm resolution. Liquid rewards were delivered automatically upon successful target acquisition and hold in the delayed-reach task, or upon successful target decode in the oBCI task, which operated in the same way but rendered a magenta annulus around the decoded target upon successful decode, or gray for unsuccessful decode.

Task timing, stimulus control, and behavior monitoring were performed using Simulink with real-time xPC target (Mathworks Inc., Natick, MA) while microscope control, image acquisition, and online decode were performed by a separate PC (Fig. 5A). Images were acquired continually throughout the duration of the task at 512 x 512 resolution using resonant scanning at 30.3 Hz frame rate using PrairieView software (Bruker Instruments, Inc.),

To construct reach-direction tuning maps, we regressed each pixel’s activity onto 2D hand velocity using a 50 ms lag between neural activity and behavior. We colored each pixel by using its X and Y coefficients to index into a perceptually uniform colormap, defined by varying luminance and chroma in the LCH color space (used in Fig. 4 H-N, Fig. 6E and Fig. 7A). To segment individual cells and processes, we identified pixels with statistically significant tuning (F-test, uncorrected for multiple comparisons, p < 0.05). We performed morphological closing followed by dilation on this thresholded image, and identified ROIs from the connected components. These geometric operations act on shapes within the image, eliminating small isolated puncta and establishing a cohesive boundary around connected neural processes. A manual inspection step identified ROIs to be merged or split. Splitting was performed using an automated approach, using k-means clustering on pixelwise tuning, followed by a geodesic distance transformation to identify cohesive new ROIs. Activity within each ROI was averaged over pixels and then over trials to produce PSTHs.

During oBCI decoding sessions, image data were accessed using PrairieLink (Bruker Instruments, Inc.), processed using a soft real-time C^++^ / Cython / NumPy pipeline, which avoided memory copies to reduce latency. The code for this real-time memory access is available at https://github.com/djoshea/obci/. Frames were ignored until a fixed period of time after the go cue (T_skip_), after which frames are integrated for a fixed duration (T_int_) (Fig. 5B). T_skip_ was selected to roughly match the time needed for visual information to reach PMd and M1, so as to avoid acquiring (for decode purposes) neural activity that is unlikely to be related to the reach target. Then, starting right after T_skip_, we acquired peri-movement neural activity during T_int_. This decoding nomenclature was used in a prior study from our group^8^. Each imaging frame was processed by applying an approximate Gaussian spatial blur (standard deviation of between 3 pixels) and accumulated into a running average (Fig. 5C). During training trials, after T_int_, the average image was accumulated into the running average for the current target, Y_i,j,k_, where k ∈ 1, 2, 3, 4 indexes target location, and i, j index pixel [x,y] location. During decode trials, after T_int_, the reach target was decoded using a pixel-wise minimum mean square error (MMSE) decoder as described in equation (1):

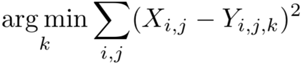

X is the image formed by averaging frames acquired during the T_int_ on the current trial. oBCI experimental sessions were often terminated to explore different imaging depths or fields of view the same day, as opposed to observing performance ultimately roll off.

### Offline analysis

Offline analysis was performed using custom MATLAB (Mathworks Inc, Natick, MA) code, available upon reasonable request.

### Assay for neutralizing antibodies

#### Serum Samples

Blood samples were collected from five rhesus macaques. Blood draws were either performed at Stanford University or at the University of Texas at Austin, depending on the location of the monkey. Approximately 1 ml of blood was drawn at each collection date. Blood samples were placed in an IEC Centra GP8 Centrifuge at 2500 RPM for 150 seconds, transferred to a 1.5 mL Eppendorf, and then centrifuged at 14000 RPM for 3 minutes in an Eppendorf Centrifuge 5430. Remaining serum was extracted and stored at -80° C until use.

#### Assay for measuring AAV neutralizing antibodies

We compared neutralizing antibody titers in blood sera collected before and after intracranial injections of AAVs using a standard *in vitro* assay (see Supplementary Table 1). HEK cells were seeded onto a clear bottom, black 96-well microplate (Corning Inc.) in dMEM/F12 without phenol red (Gibco Inc.) containing 10% Fetal Bovin Serum and incubated for 24 hours at 37° C with 5% CO_2_. The next day, serum dilutions were prepared (1:5, 1:25, 1:50 and 1:250), added to diluted adeno-associated virus encoding eYFP (AAV-Ef1α-eYFP), 1:1 volume-to-volume ratio, and for 1 hour at 37 °C (see Supplementary Table 2 for AAV concentrations). Next, 10 μl of the serum-virus mixture was added to each well of cultured cells, resulting in final serum concentrations or 1:100, 1:500, 1:1000 and 1:5000. Experiments were run in triplicate for each condition. Positive control (no serum) and negative control (no AAV) wells were included in each experiment. For the pre-injection neutralization experiment, two serum dilutions were prepared (1:5 and 1:25), added to diluted adeno-associated virus encoding eYFP (AAV-Ef1α-eYFP), 1:1 volume-to-volume ratio, and for 1 hour at 37 °C (see Table 2 for AAV concentrations). Next, 10 μl of the serum-virus mixture was added to 6 wells of cultured cells. A positive, no serum, control was included for each serotype.

#### Fluorescence quantification

Thirty-six or forty-eight hours after incubation, wells were imaged and YFP fluorescence measured. Fluorescence was quantified using a Tecan Infinite M1000 Microplate Reader. For each serum dilution, pre- and post-injection sera were compared using unpaired t tests. In addition, widefield images of each well were captured using a Leica DMi8 microscope (10x objective) to visualize YFP expression.

### CLARITY tissue clearing and imaging

The area of interest was dissected from the whole brain and embedded in 1% hydrogel. The tissue was cleared in the SmartClear (Lifecanvas Inc.) for 2 weeks, then stained with anti-GFP conjugated to alexa-647 (Invitrogen Inc.) and lectin conjugated to DyLight 488 (antibody concentrations 1:100 in PBST with 1% triton-x & 0.2% sodium azide) for one week, and finally washed in PBST for 1 week prior to imaging. The tissue was immersed in RapiClear (SunJin Lab Co) for 2 days and then imaged using an Olympus two-photon microscope with 10x 0.6 NA CLARITY objective.

## Author contributions

E.M.T and D.J.O designed and constructed the 2P imaging system and designed the implant. E.M.T, D.J.O, X.S. performed virus injections, designed and conducted the *in-vivo* imaging experiments. D.J.O. implemented the real-time decoding system. E.M.T, D.J.O, and G.B. performed analysis of 2P imaging. S.I.R., E.M.T, and D.J.O performed the surgeries. J.H.M. provided extensive assistance with all aspects of imaging, experimental design, analysis, and manuscript preparation. A.C. performed CLARITY imaging. B.H. performed histology and CLARITY sample preparation. S.V. performed neutralizing antibody assay and provided assistance with manuscript revision. L.C. provided assistance with implant and equipment design and testing. G.B. developed 2P imaging analysis code. W.A. and I.K. built the widefield imaging microscope and provided significant assistance with 2P imaging. S.Q. provided assistance with 2P imaging. M.M. performed surgery and assistance with virus injections in Monkey X. Y.C., M.W., and E.S., provided assistance with implant design and performed widefield imaging and GCaMP validation in Monkey L. C.R. provided assistance selecting and validating viral constructs. M.S. provided assistance with 2P analysis. K.D. and K.V.S. provided guidance and assistance with all aspects of the work. E.M.T., D.J.O., X.S., and S.V. wrote the manuscript with input and editing from all authors.

## Code availability

Offline analysis was performed using custom MATLAB (Mathworks Inc., Natick, MA) code, available upon reasonable request. The code for the real-time decoder is available at https://github.com/djoshea/obci.

## Data Availability

The datasets generated during and/or analyzed during the current study are available from the corresponding authors upon reasonable request.

## Acknowledgements

We thank M. Risch, M. Wechsler, L. Yates and R. Reeder for surgical assistance and veterinary care and B. Davis for administrative support. This work was supported by NIH NRSA grant 1F31NS089376-01 (E.M.T.), a Stanford Graduate Fellowship (E.M.T.), NSF IGERT grant 0734683 (E.M.T.), DARPA BTO ‘‘NeuroFAST’’ award W911NF-14-2-0013 (K.D. and K.V.S.), the Howard Hughes Medical Institute (K.D. and K.V.S.), an NIH Director’s Pioneer Award 8DP1HD075623, and the Simons Foundation Collaboration on the Global Brain awards 325380 and 543045 (K.V.S.). We thank John Rafter, Aaron Statz, and Michael J. Fox for their assistance with the 2P imaging microscope and achieving real-time access to microscope data.

## Declaration of interests

K.V.S. is a consultant to Neuralink Corp. and on the Scientific Advisory Boards of CTRL-Labs Inc., Mind-X Inc., Inscopix Inc. and Heal Inc. K.D. is on the scientific advisory board of Circuit Therapeutics. These entities did not support this work. Following this study, J.H.M is now a member of the scientific advisory board of Bruker.

## Supplementary Figures

**Supplementary Figure S1.**
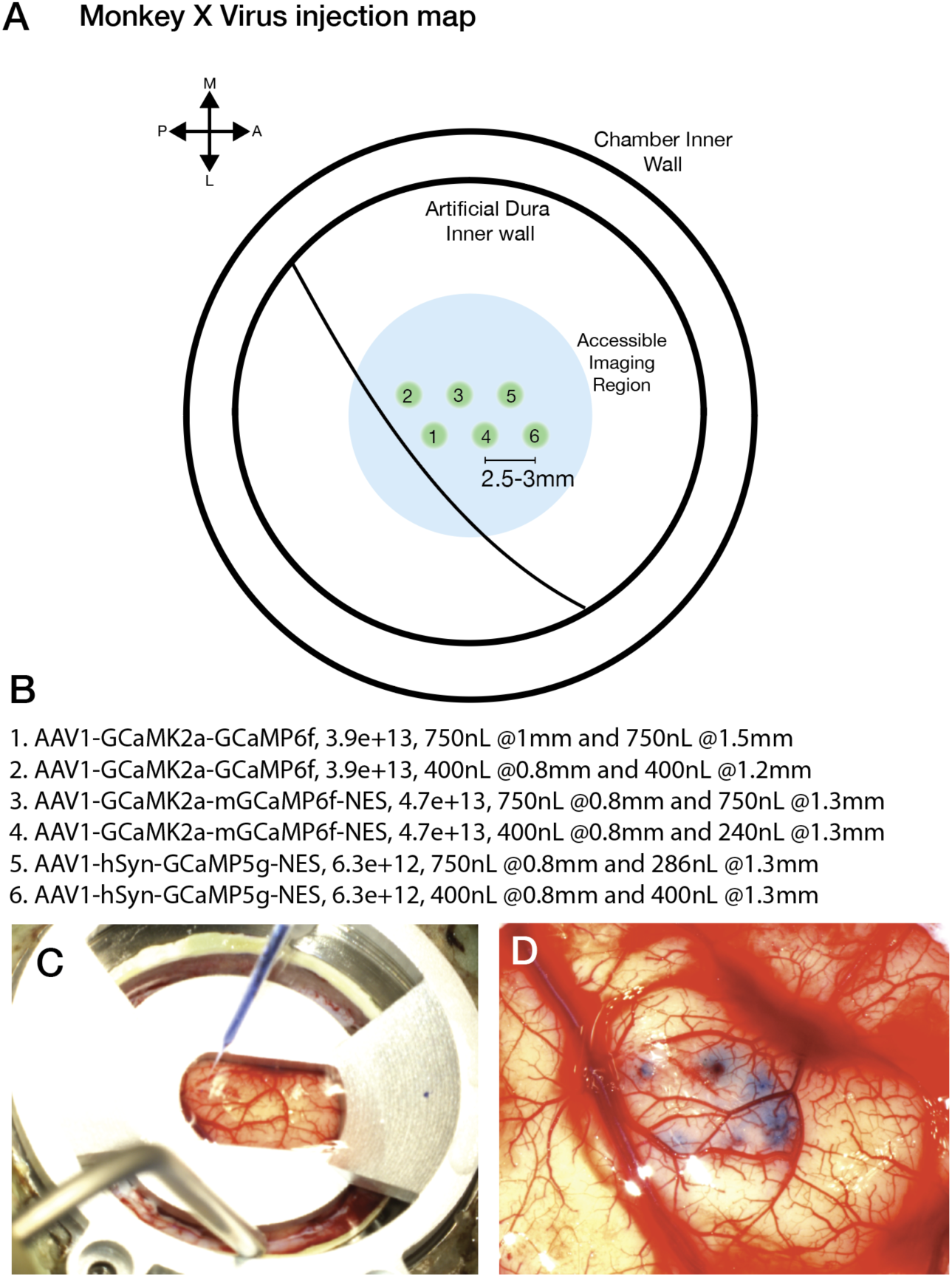
Virus injection map and cortical surface pictures. (**A**) Viral construct injection map for monkey X. (**B**) List of injected viral constructs and injection volumes and concentrations (**C**) Tissue stabilization apparatus and glass pipette during injections (**D**) Injection sites from (**A**), visible via Trypan blue dye mixed with viral constructs.

**Supplementary Figure S2.**
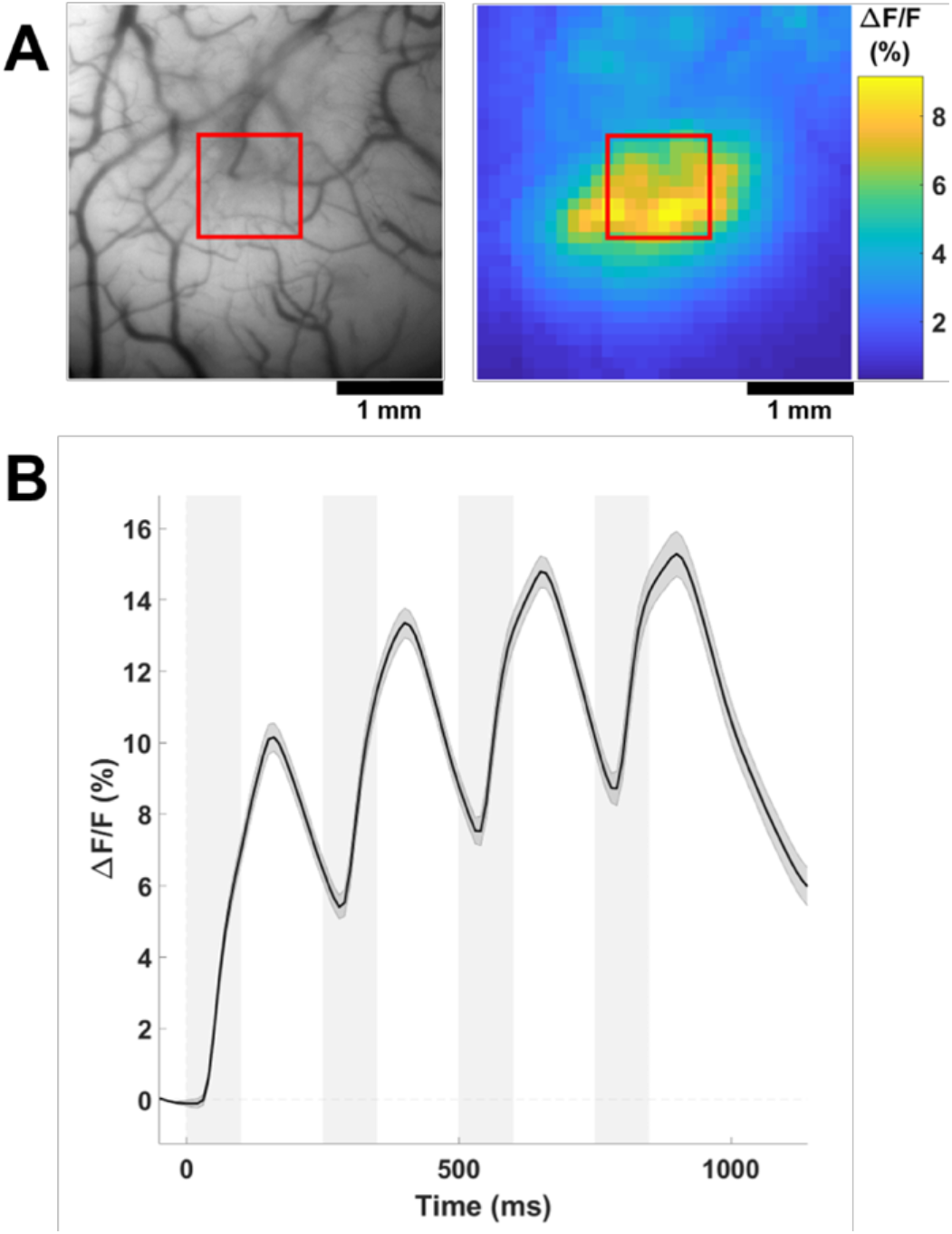
Reliable widefield GCaMP signal from macaque visual cortex. **(A)** Reference vasculature (left) and widefield signal in response to 4Hz flashed grating (right) at the intersection of two sites injected with AAV1-CaMKIIa-NES-GCaMP6f. In the response map, color indicates amplitude of the 4 Hz FFT component computed at each location. **(B)** Average time course of GCaMP response to a 4 Hz flashed grating (100 ms on, 150 ms off) with stimulus presentation time shaded in gray. Shaded area around the time course signal represents ± SEM over 10 stimulus repetitions. Note that the GCaMP signal does not quickly return to baseline after each stimulus presentation, so a buildup in the response can be seen over time, as reported previously with CaMKIIα-GCaMP^18^. The 1x1 mm ROI used for the time course is indicated by red squares in **(A)**. Example signal was recorded 14 weeks post-injection and continues to provide reliable signal after 13 months.

**Supplementary Figure S3.**
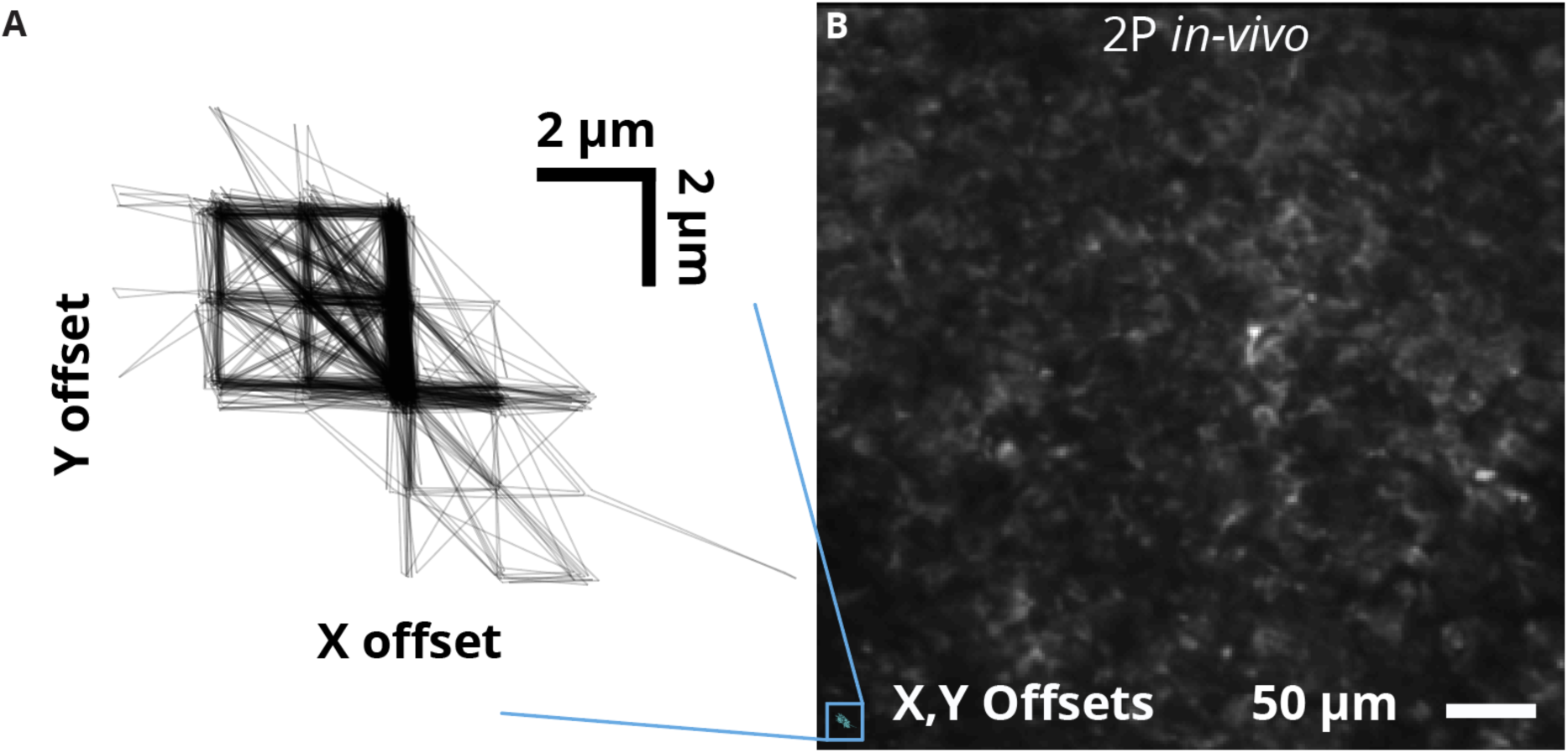
Image stability during motor behavior. (**A)** X and Y pixel offsets after image registration. The majority of frames require only one or two pixels worth of shift, illustrating the high degree of image stability during behavior. (**B)** Mean image from a single time series, with overlay of pixel offsets from A) in bottom left corner.

**Supplementary Figure S4.**
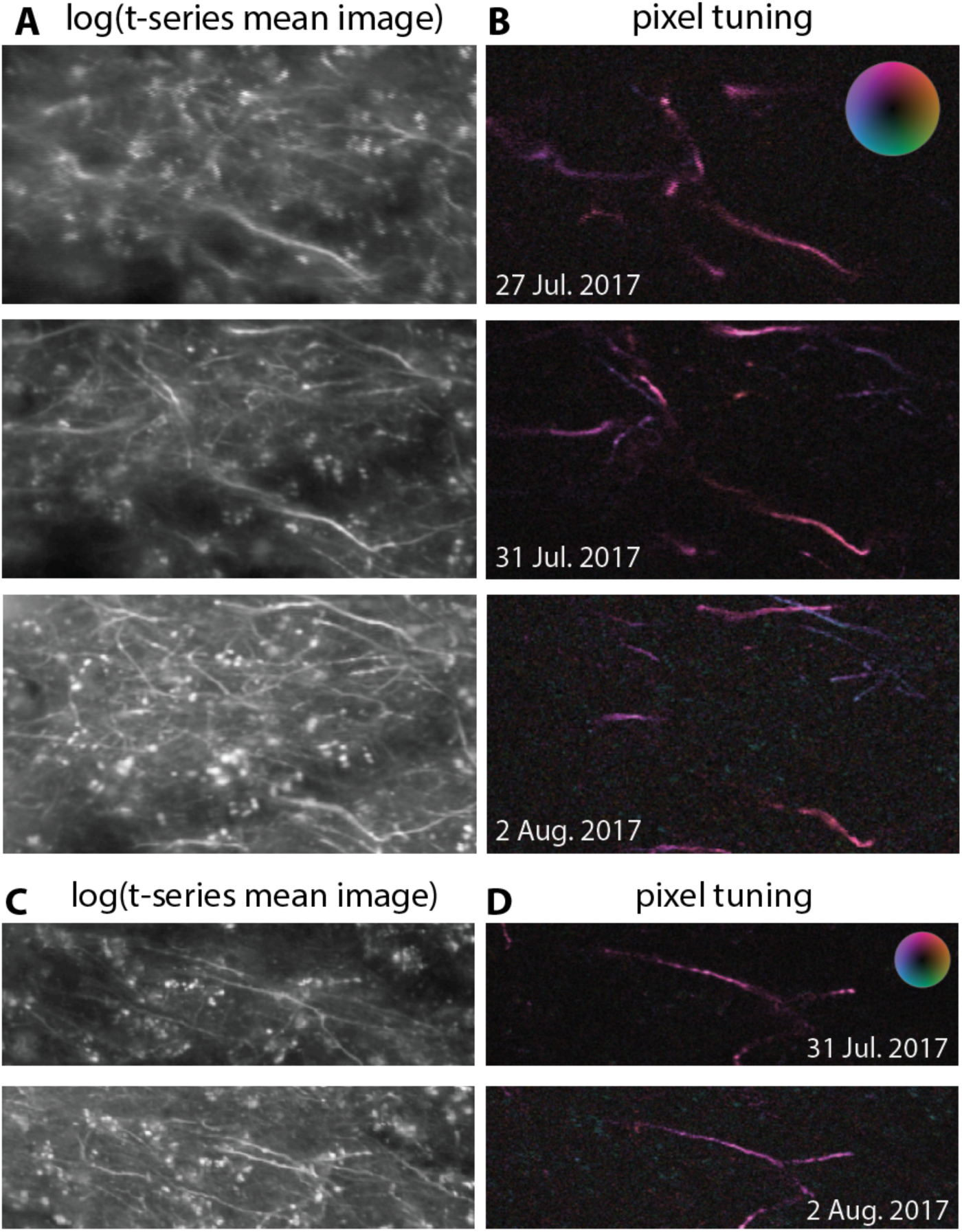
Additional examples of consistent neural tuning observed across multiple sessions. **(A)** The same neural process observed during four-target reaching behavior is shown across three experimental sessions spanning six. Time series data were aligned offline and pixel-wise tuning was compute by regressing the raw pixel values against arm velocity. **(B)** The mean of each time series was computed, and log transformed to highlight dim features for visualization purposes. **(C-D)** An additional example of stable tuning across multiple experimental sessions.

**Supplementary Figure S5.**
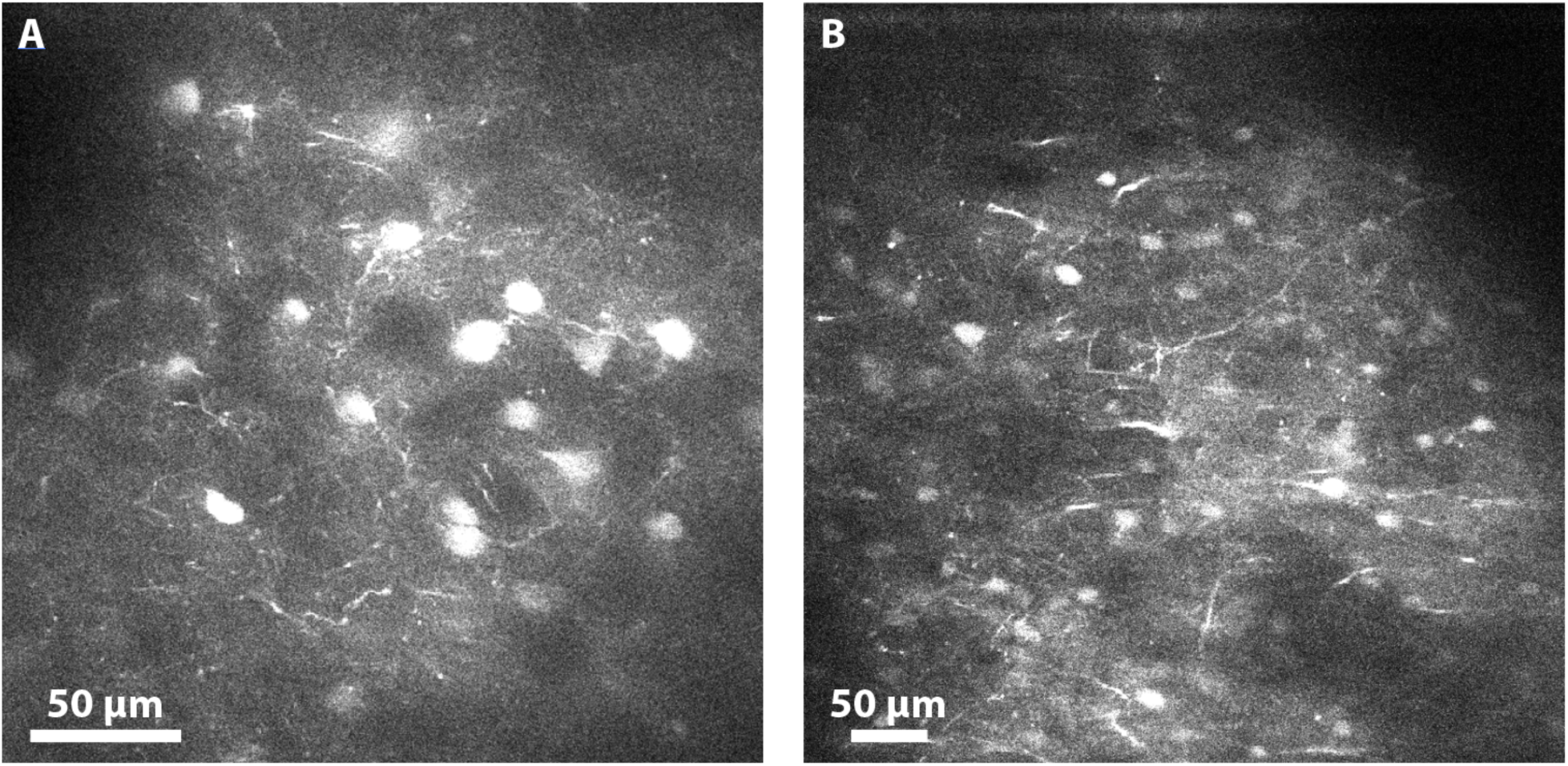
Cells exhibiting non-standard GCaMP expression. **(A,B)** Two fields of view from Monkey W showing numerous brightly fluorescent cells that do not exhibit a “halo” pattern characteristic of typical calcium imaging. These cells do not modulate in fluorescence during behavior. We observed this phenomen at multiple injection sites and multiple subjects and a range of viral serotypes, including AAV1 and AAV with CAMKIIa and hSyn promoters. While we would suspect unhealthy overexpression of GCaMP due to filled nuclei and lack of traditional “halo” shape, we observed the presence of these cells within six weeks of injecting virus. In addition, such cells did not exhibit healthy levels of fluorescence before displaying bright, filled nuclei. Instead, they exhibit no fluorescence and appeared as shown within several days. This suggests that overexpression may not be the primary explanation for this phenomenon. Scale bars are approximate.

## Supplementary Tables

**Table 1.**
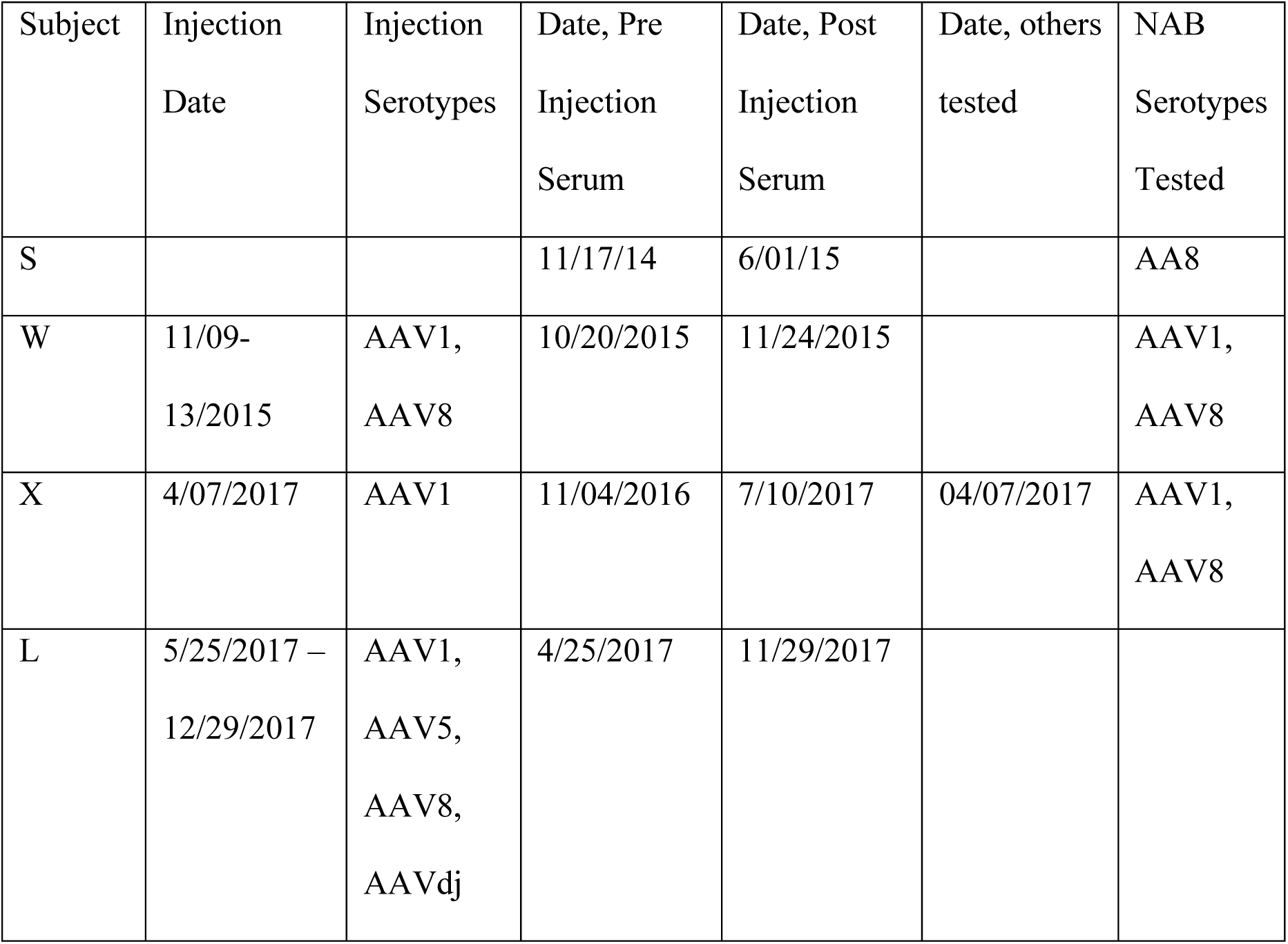

| Subject | Injection<br>Date | Injection<br>Serotypes | Date, Pre<br>Injection<br>Serum | Date, Post<br>Injection<br>Serum | Date, others<br>tested | NAB<br>Serotypes<br>Tested |
| --- | --- | --- | --- | --- | --- | --- |
| S |  |  | 11/17/14 | 6/01/15 |  | AA8 |
| W | 11/09-13/2015 | AAV1,<br>AAV8 | 10/20/2015 | 11/24/2015 |  | AAV1,<br>AAV8 |
| X | 4/07/2017 | AAV1 | 11/04/2016 | 7/10/2017 | 04/07/2017 | AAV1,<br>AAV8 |
| L | 5/25/2017 –<br>12/29/2017 | AAV1,<br>AAV5,<br>AAV8,<br>AAVdj | 4/25/2017 | 11/29/2017 |  |  |

**Table 2.**

| Virus | Original Titer | Diluted titer used |
| --- | --- | --- |
| AAV1-Ef1a-YFP | 1.8E13 | 1.8E11 |
| AAV8-Ef1a-eYFP | 2.4E14 | 2.4E12 |

## Notes

#### Summary of Updates

- added middle initial for one author - removed extraneous line from one figure panel

## References

1. Collinger, J. L. et al. High-performance neuroprosthetic control by an individual with tetraplegia. Lancet 381, 557–564 (2013).

2. Pandarinath, C. et al. Neural population dynamics in human motor cortex during movements in people with ALS. eLife 4, e07436 (2015).

3. Shenoy, K. V. & Carmena, J. M. Combining Decoder Design and Neural Adaptation in Brain-Machine Interfaces. Neuron 84, 665–680 (2014).

4. Taylor, D. M., Tillery, S. I. H. & Schwartz, A. B. Direct cortical control of 3D neuroprosthetic devices. Science 296, 1829–1832 (2002).

5. Serruya, M. D., Hatsopoulos, N. G., Paninski, L., Fellows, M. R. & Donoghue, J. P. Instant neural control of a movement signal. Nature 416, 141–142 (2002).

6. Carmena, J. M. et al. Learning to control a brain-machine interface for reaching and grasping by primates. PLoS biology 1, 193–208 (2003).

7. Musallam, S., Corneil, B. D., Greger, B., Scherberger, H. & Andersen, R. A. Cognitive control signals for neural prosthetics. Science 305, 258–262 (2004).

8. Santhanam, G., Ryu, S. I., Yu, B. M., Afshar, A. & Shenoy, K. V. A high-performance brain– computer interface. Nature 442, 195–198 (2006).

9. Moritz, C. T., Perlmutter, S. I. & Fetz, E. E. Direct control of paralysed muscles by cortical neurons. Nature 456, 639–642 (2008).

10. Velliste, M., Perel, S., Spalding, M. C., Whitford, A. S. & Schwartz, A. B. Cortical control of a prosthetic arm for self-feeding. Nature 453, 1098–1101 (2008).

11. Ethier, C., Oby, E. R., Bauman, M. J. & Miller, L. E. Restoration of grasp following paralysis through brain-controlled stimulation of muscles. 485, (Nature Publishing Group, 2012).

12. Gilja, V. et al. A high-performance neural prosthesis enabled by control algorithm design. Nature Neuroscience 15, 1752–1757 (2012).

13. Kao, J. C., Nuyujukian, P., Ryu, S. I., Churchland, M. M. & Cunningham, J. P. Single-trial dynamics of motor cortex and their applications to brain-machine interfaces. Nature Communications 6, 1–12 (2015).

14. Shenoy, K. V., Sahani, M. & Churchland, M. M. Cortical control of arm movements: a dynamical systems perspective. Annu. Rev. Neurosci. 36, 337–359 (2013).

15. Heider, B., Nathanson, J. L., Isacoff, E. Y., Callaway, E. M. & Siegel, R. M. Two-photon imaging of calcium in virally transfected striate cortical neurons of behaving monkey. PLoS ONE 5, 1–13 (2010).

16. Ju, N., Jiang, R., Macknik, S. L., Martinez-Conde, S. & Tang, S. Long-term all-optical interrogation of cortical neurons in awake-behaving nonhuman primates. PLOS Biology 16, e2005839 (2018).

17. Li, M., Liu, F., Jiang, H., Lee, T. S. & Tang, S. Long-Term Two-Photon Imaging in Awake Macaque Monkey. Neuron 93, 1049–1057.e3 (2017).

18. Seidemann, E. et al. Calcium imaging with genetically encoded indicators in behaving primates. Elife 5, (2016).

19. Garg, A. K., Li, P., Rashid, M. S. & Callaway, E. M. Color and orientation are jointly coded and spatially organized in primate primary visual cortex. *Science (New York*, N.Y.) 364, 1275–1279 (2019).

20. Sadakane, O. et al. Long-Term Two-Photon Calcium Imaging of Neuronal Populations with Subcellular Resolution in Adult Non-human Primates. Cell Reports 13, 1989–1999 (2015).

21. Yamada, Y., Matsumoto, Y., Okahara, N. & Mikoshiba, K. Chronic multiscale imaging of neuronal activity in the awake common marmoset. Scientific Reports 6, 35722 (2016).

22. Ebina, T. et al. Two-photon imaging of neuronal activity in motor cortex of marmosets during upper-limb movement tasks. Nature Communications 9, 1–16 (2018).

23. Sofroniew, N. J., Flickinger, D., King, J. & Svoboda, K. A large field of view two-photon mesoscope with subcellular resolution for in vivo imaging. Elife 5, (2016).

24. Stringer, C., Pachitariu, M., Steinmetz, N., Carandini, M. & Harris, K. D. High-dimensional geometry of population responses in visual cortex. Nature 571, 361–365 (2019).

25. Trautmann, E. M. et al. Accurate Estimation of Neural Population Dynamics without Spike Sorting. Neuron 103, 292–308.e4 (2019).

26. Jung, J. C., Mehta, A. D., Aksay, E., Stepnoski, R. & Schnitzer, M. J. In vivo mammalian brain imaging using one- and two-photon fluorescence microendoscopy. J. Neurophysiol. 92, 3121–3133 (2004).

27. Bollimunta, A., et al. Head mounted microendoscopic calcium imaging in deep layer premotor cortex of behaving rhesus macaque. in (2019).

28. Trautmann, E. et al. Spatially heterogenous tuning in rhesus motor cortex revealed using neuropixels probes. in Society for Neuroscience (2019).

29. Peters, A. J., Lee, J., Hedrick, N. G., O’Neil, K. & Komiyama, T. Reorganization of corticospinal output during motor learning. Nature Neuroscience 20, 1133–1141 (2017).

30. Ranganathan, G. N. et al. Active dendritic integration and mixed neocortical network representations during an adaptive sensing behavior. Nat Neurosci 21, 1583–1590 (2018).

31. Takahashi, N., Oertner, T. G., Hegemann, P. & Larkum, M. E. Active cortical dendrites modulate perception. Science 354, 1587–1590 (2016).

32. Xu, N. et al. Nonlinear dendritic integration of sensory and motor input during an active sensing task. Nature 492, 247–251 (2012).

33. Beaulieu-Laroche, L., Toloza, E. H. S., Brown, N. J. & Harnett, M. T. Widespread and Highly Correlated Somato-dendritic Activity in Cortical Layer 5 Neurons. Neuron 103, 235–241.e4 (2019).

34. Chung, K. & Deisseroth, K. CLARITY for mapping the nervous system. Nature methods 10, 508–13 (2013).

35. Tomer, R., Ye, L., Hsueh, B. & Deisseroth, K. Advanced CLARITY for rapid and high-resolution imaging of intact tissues. Nature protocols 9, 1682–97 (2014).

36. O’Shea, D. J. et al. The need for calcium imaging in nonhuman primates: New motor neuroscience and brain-machine interfaces. Experimental Neurology 287, 437–451 (2017).

37. Sadtler, P. T. et al. Neural constraints on learning. Nature 512, 423–426 (2014).

38. Peixoto, D. et al. Population dynamics of choice representation in dorsal premotor and primary motor cortex. bioRxiv 283960 (2018). doi:10.1101/283960

39. Arieli, A., Grinvald, A. & Slovin, H. Dural substitute for long-term imaging of cortical activity in behaving monkeys and its clinical implications. Journal of Neuroscience Methods 114, 119–133 (2002).

40. Shtoyerman, E., Arieli, A., Slovin, H., Vanzetta, I. & Grinvald, A. Long-term optical imaging and spectroscopy reveal mechanisms underlying the intrinsic signal and stability of cortical maps in V1 of behaving monkeys. Journal of Neuroscience 20, 8111–8121 (2000).

41. Davis, T. S., Torab, K., House, P. & Greger, B. A minimally invasive approach to long-term head fixation in behaving nonhuman primates. Journal of Neuroscience Methods 181, 106–110 (2009).

42. Azimi, K., Prescott, I. A., Marino, R. A., Winterborn, A. & Levy, R. Low profile halo head fixation in non-human primates. Journal of Neuroscience Methods 268, 23–30 (2016).

43. Isoda, M. et al. Design of a head fixation device for experiments in behaving monkeys. Journal of Neuroscience Methods 141, 277–282 (2005).

44. Kotterman, M. A. et al. Antibody neutralization poses a barrier to intravitreal adeno-associated viral vector gene delivery to non-human primates. Gene Therapy 22, 116–126 (2014).

45. Mingozzi, F. & High, K. A. Immune responses to AAV vectors: overcoming barriers to successful gene therapy. Blood 122, 23–36 (2013).

46. Calcedo, R. et al. Preexisting Neutralizing Antibodies to Adeno-Associated Virus Capsids in Large Animals Other Than Monkeys May Confound In Vivo Gene Therapy Studies. Human gene therapy methods 26, 103–105 (2015).

47. Mendoza, S. D., El-Shamayleh, Y. & Horwitz, G. D. AAV-mediated delivery of optogenetic constructs to the macaque brain triggers humoral immune responses. Journal of Neurophysiology 117, 2004–2013 (2017).

48. Kaufman, M. T. et al. The Largest Response Component in the Motor Cortex Reflects Movement Timing but Not Movement Type. eNeuro 3, (2016).

49. Cunningham, J. P. & Yu, B. M. Dimensionality reduction for large-scale neural recordings. Nat. Neurosci. 17, 1500–1509 (2014).

50. Yu, B. M. et al. Gaussian-Process Factor Analysis for Low-Dimensional Single-Trial Analysis of Neural Population Activity. Journal of Neurophysiology 614 – 635 (2009). doi:10.1152/jn.90941.2008.

51. Shenoy, K. V. et al. Neural prosthetic control signals from plan activity. Neuroreport 14, 591–596 (2003).

52. Kandel, E. R., Schwartz, J. H., Jessell, T. M., Siegelbaum, S. A. & Hudspeth, A. J. Principles of Neural Science, Fifth Edition. (McGraw-Hill Education, 2012).

53. Kalaska, J. F. Emerging ideas and tools to study the emergent properties of the cortical neural circuits for voluntary motor control in non-human primates. F1000Res 8, (2019).

54. Jun, J. J. et al. Fully integrated silicon probes for high-density recording of neural activity. Nature 551, 232–236 (2017).

55. Dadarlat, M. C., O’Doherty, J. E. & Sabes, P. N. A learning-based approach to artificial sensory feedback leads to optimal integration. Nat. Neurosci. 18, 138–144 (2015).

56. Flesher, S. N. et al. Intracortical microstimulation of human somatosensory cortex. Sci. Transl. Med. 8, 361ra141 (2016).

57. George, J. A. et al. Biomimetic sensory feedback through peripheral nerve stimulation improves dexterous use of a bionic hand. Science Robotics 4, eaax2352 (2019).

58. Histed, M. H., Bonin, V. & Reid, R. C. Direct Activation of Sparse, Distributed Populations of Cortical Neurons by Electrical Microstimulation. Neuron 63, 508–522 (2009).

59. O’Shea, D. J. & Shenoy, K. V. ERAASR: an algorithm for removing electrical stimulation artifacts from multielectrode array recordings. J. Neural Eng. 15, 026020 (2018).

60. Dana, H. et al. Sensitive red protein calcium indicators for imaging neural activity. eLife 5, e12727 (2016).

61. Clancy, K. B., Koralek, A. C., Costa, R. M., Feldman, D. E. & Carmena, J. M. Volitional modulation of optically recorded calcium signals during neuroprosthetic learning. Nature neuroscience 17, 807– 809 (2014).

62. Vyas, S. et al. Neural Population Dynamics Underlying Motor Learning Transfer. Neuron 97, 1177– 1186.e3 (2018).

63. Grewe, B. F. & Helmchen, F. Optical probing of neuronal ensemble activity. Current opinion in neurobiology 19, 520–9 (2009).

64. Peron, S., Chen, T.-W. & Svoboda, K. Comprehensive imaging of cortical networks. Current Opinion in Neurobiology 32, 115–123 (2015).

65. Barthó, P. et al. Characterization of neocortical principal cells and interneurons by network interactions and extracellular features. Journal of neurophysiology 92, 600–608 (2004).

66. Kaufman, M. T. et al. Roles of monkey premotor neuron classes in movement preparation and execution. Journal of neurophysiology 104, 799–810 (2010).

67. Kaufman, M. T., Churchland, M. M. & Shenoy, K. V. The roles of monkey M1 neuron classes in movement preparation and execution. Journal of Neurophysiology 110, 817–825 (2013).

68. Jia, X. et al. High-density extracellular probes reveal dendritic backpropagation and facilitate neuron classification. Journal of Neurophysiology 121, 1831–1847 (2019).

69. Steinmetz, N. A., Koch, C., Harris, K. D. & Carandini, M. Challenges and opportunities for large-scale electrophysiology with Neuropixels probes. Current Opinion in Neurobiology 50, 92–100 (2018).

70. Krienen, F. M. et al. Innovations in Primate Interneuron Repertoire. bioRxiv 709501 (2019). doi:10.1101/709501

71. Tasic, B. et al. Shared and distinct transcriptomic cell types across neocortical areas. Nature 563, 72– 78 (2018).

72. Wang, X. et al. Three-dimensional intact-tissue sequencing of single-cell transcriptional states. Science 361, eaat5691 (2018).

73. Hsueh, B. et al. Pathways to clinical CLARITY: volumetric analysis of irregular, soft, and heterogeneous tissues in development and disease. Sci Rep 7, 1–16 (2017).

74. Economo, M. N. et al. Distinct descending motor cortex pathways and their roles in movement. Nature 563, 79–84 (2018).

75. Li, N., Chen, T., Guo, Z. V., Gerfen, C. R. & Svoboda, K. A motor cortex circuit for motor planning and movement. Nature 519, 51–56 (2015).

76. Barretto, R. P. J., Messerschmidt, B. & Schnitzer, M. J. In vivo fluorescence imaging with high-resolution microlenses. Nature Methods 6, 511–512 (2009).

77. Abdelfattah, A. S. et al. Bright and photostable chemigenetic indicators for extended in vivo voltage imaging. Science eaav6416 (2019). doi:10.1126/science.aav6416

78. Anderson, H. E., Fontaine, A. K., Caldwell, J. H. & Weir, R. F. Imaging of electrical activity in small diameter fibers of the murine peripheral nerve with virally-delivered GCaMP6f. Sci Rep 8, 1–9 (2018).

79. Gunaydin, L. A. et al. Natural Neural Projection Dynamics Underlying Social Behavior. Cell 157, 1535–1551 (2014).

80. Lewis, D. A. et al. Dopamine transporter immunoreactivity in monkey cerebral cortex: regional, laminar, and ultrastructural localization. J. Comp. Neurol. 432, 119–136 (2001).

81. Brombas, A., Fletcher, L. N. & Williams, S. R. Activity-dependent modulation of layer 1 inhibitory neocortical circuits by acetylcholine. J. Neurosci. 34, 1932–1941 (2014).

82. Herrero, J. L., Gieselmann, M. A. & Thiele, A. Muscarinic and Nicotinic Contribution to Contrast Sensitivity of Macaque Area V1 Neurons. Front. Neural Circuits 11, (2017).

83. Soma, S., Shimegi, S., Osaki, H. & Sato, H. Cholinergic modulation of response gain in the primary visual cortex of the macaque. J. Neurophysiol. 107, 283–291 (2012).

84. Croxson, P. L., Kyriazis, D. A. & Baxter, M. G. Cholinergic modulation of a specific memory function of prefrontal cortex. Nature Neuroscience 14, 1510–1512 (2011).

85. Saunders, A. et al. A direct GABAergic output from the basal ganglia to frontal cortex. Nature 521, 85–89 (2015).

86. Strick, P. L. & Sterling, P. Synaptic termination of afferents from the ventrolateral nucleus of the thalamus in the cat motor cortex. A light and electron microscopy study. J. Comp. Neurol. 153, 77– 106 (1974).

87. Roe, A. W., Chernov, M. M., Friedman, R. M. & Chen, G. In Vivo Mapping of Cortical Columnar Networks in the Monkey with Focal Electrical and Optical Stimulation. Front Neuroanat 9, 135 (2015).

88. Perich, M. G., Gallego, J. A. & Miller, L. E. A Neural Population Mechanism for Rapid Learning. Neuron 100, 964–976.e7 (2018).

89. Golub, M. D. et al. Learning by neural reassociation. Nat. Neurosci. 21, 607–616 (2018).

90. Oby, E. R. et al. New neural activity patterns emerge with long-term learning. Proc. Natl. Acad. Sci. U.S.A. 116, 15210–15215 (2019).

91. Orsborn, A. L. et al. Closed-Loop Decoder Adaptation Shapes Neural Plasticity for Skillful Neuroprosthetic Control. Neuron 82, 1380–1393 (2014).

92. Green, A. M. & Kalaska, J. F. Learning to move machines with the mind. Trends in Neurosciences 34, 61–75 (2011).

93. Ganguly, K. & Carmena, J. M. Emergence of a stable cortical map for neuroprosthetic control. PLoS Biol. 7, e1000153 (2009).

94. Santhanam, G. et al. HermesB: a continuous neural recording system for freely behaving primates. IEEE Trans. Biomed. Eng. 54, 2037–2050 (2007).

95. Tolias, A. S. et al. Recording chronically from the same neurons in awake, behaving primates. J. Neurophysiol. 98, 3780–3790 (2007).

96. Stevenson, I. H. et al. Statistical assessment of the stability of neural movement representations. Journal of neurophysiology 106, 764–774 (2011).

97. Fraser, G. W. & Schwartz, A. B. Recording from the same neurons chronically in motor cortex. Journal of Neurophysiology 107, 1970–1978 (2012).

98. Chen, S. X., Kim, A. N., Peters, A. J. & Komiyama, T. Subtype-specific plasticity of inhibitory circuits in motor cortex during motor learning. Nature Neuroscience 18, 1109–1115 (2015).

99. Driscoll, L. N., Pettit, N. L., Minderer, M., Chettih, S. N. & Harvey, C. D. Dynamic Reorganization of Neuronal Activity Patterns in Parietal Cortex. Cell 170, 986–999.e16 (2017).

100. Huber, D., Gutnisky, D. & Peron, S. Multiple dynamic representations in the motor cortex during sensorimotor learning. Nature 484, 473–478 (2012).

101. Margolis, D. J. et al. Reorganization of cortical population activity imaged throughout long-term sensory deprivation. Nature Neuroscience 15, 1539–1546 (2012).

102. Peters, A. J., Chen, S. X. & Komiyama, T. Emergence of reproducible spatiotemporal activity during motor learning. Nature 510, 263–267 (2014).

103. Ziv, Y. et al. Long-term dynamics of CA1 hippocampal place codes. Nature neuroscience 16, 264– 6 (2013).

104. Sun, X., Kao, J. C., Marshel, J. H., Ryu, S. I. & Shenoy, K. V. Feasibility analysis of genetically-encoded calcium indicators as a neural signal source for all-optical brain-machine interfaces. in 2017 8th International IEEE/EMBS Conference on Neural Engineering (NER) 174–180 (IEEE, 2017).

105. Helassa, N., Podor, B., Fine, A. & Török, K. Design and mechanistic insight into ultrafast calcium indicators for monitoring intracellular calcium dynamics. Sci Rep 6, (2016).

106. Deisseroth, K. & Schnitzer, M. J. Engineering approaches to illuminating brain structure and dynamics. Neuron 80, 568–77 (2013).

107. Chen, T.-W. et al. Ultrasensitive fluorescent proteins for imaging neuronal activity. Nature 499, 295–300 (2013).

108. Emiliani, V., Cohen, A. E., Deisseroth, K. & Hausser, M. All-Optical Interrogation of Neural Circuits. Journal of Neuroscience 35, 13917–13926 (2015).

109. Tang, S. et al. Complex Pattern Selectivity in Macaque Primary Visual Cortex Revealed by Large-Scale Two-Photon Imaging. Current Biology 28, 38–48.e3 (2018).

110. Galvan, A. et al. Nonhuman Primate Optogenetics: Recent Advances and Future Directions. J. Neurosci. 37, 10894–10903 (2017).

111. Watakabe, A. et al. Comparative analyses of adeno-associated viral vector serotypes 1, 2, 5, 8 and 9 in marmoset, mouse and macaque cerebral cortex. Neuroscience Research 93, 144–157 (2015).

112. Churchland, M. M., Afshar, A. & Shenoy, K. V. A central source of movement variability. Neuron 52, 1085–1096 (2006).

